# Erythropoietin Mediates Glycerophospholipid Remodeling During Human Early Erythropoiesis

**DOI:** 10.64898/2026.02.20.707106

**Authors:** Natascha Schippel, Jing Wei, Xiaokuang Ma, Jinhua Chi, Haiwei Gu, Shenfeng Qiu, Peter Stoilov, Shalini Sharma

## Abstract

The unique biconcave morphology and deformability of mature red blood cells (RBCs) require a specific membrane lipid composition. Disruption of this special lipid profile is seen in multiple types of anemia and bone marrow (BM) failure diseases, yet molecular mechanisms regulating lipid metabolism during normal erythroid differentiation remain poorly defined. Here, we identify a previously undescribed role for erythropoietin (Epo) in contributing to the appropriate lipid composition of differentiating human erythroid cells. Using single-cell transcriptomic profiling of *ex vivo* cultures of human BM-derived hematopoietic stem and progenitor cells cultured with or without Epo, we delineated transcriptional dynamics across differentiation stages, identifying the Epo-dependent transition of burst-forming unit erythroid (BFU-E) to colony-forming unit erythroid (CFU-E) progenitors. Comparative analysis revealed the activation of canonical erythroid programs involving heme biosynthesis, globin expression, and iron regulation, and additionally, uncovered a transient upregulation of lipid metabolic pathways during the BFU-E to CFU-E transition. Complementary untargeted lipidomics demonstrated Epo-dependent alterations in specific glycerophospholipid (GPL) species, consistent with differential expression of GPL biosynthesis genes in the single cell dataset. Intracellular flow cytometry further confirmed the requirement of Epo for maintaining enzymes critical for phosphatidylcholine and phosphatidylethanolamine synthesis in erythroid cells. Together, these multiomic findings reveal a new role for Epo in modulating lipid metabolism during early erythropoiesis and provide mechanistic insight into how membrane lipid composition is dynamically regulated to support normal red cell development.

## Introduction

Mature red blood cells (RBCs) are the most abundant cell type in peripheral blood and maintenance of their levels requires daily production of ∼0.2 trillion cells, or roughly two million cells per second [1–3]. The unique biconcave disc shape and deformability of RBCs are critical for their function, allowing passage through narrow capillaries while still maintaining large surface area for oxygen delivery. This characteristic morphology is partly supported by a specific membrane lipid composition, consisting of ∼65% glycerophospholipids (GPLs), including sphingolipids (SL), ∼25% cholesterol, and ∼10% glycolipids [4,5]. A key feature of the RBC membrane is the asymmetric distribution of GPLs across the lipid bilayer: phosphatidylserine (PS) and phosphatidylethanolamine (PE) are predominantly localized to the inner leaflet, whereas phosphatidylcholine (PC) and SLs are enriched in the outer leaflet [6,7]. This composition and asymmetric distribution play an important role in maintaining RBC membrane flexibility, mechanical stability, and lifespan [8–13].

Abnormal membrane lipid composition and organization are seen in anemias and other chronic diseases that are associated with reduced production and lysis of RBCs [14–17]. In high phosphatidyl choline hemolytic anemia, an accumulation of PC and cholesterol is seen in RBC membranes in the background of normal serum levels and has been linked to cell dehydration and lysis [18–21]. Elevated RBC PC levels have also been reported in autoimmune hemolytic anemia and hereditary stomatocytosis [22,23]. In hereditary hemolytic anemia, an increase in the PC to PE ratio was shown to affect membrane permeability and dysregulated ion equilibrium, resulting in hemolysis [16,20]. Anemia secondary to liver disease is associated with an increased cholesterol/GPL ratio, also reducing RBC deformability and increasing hemolysis [24]. Imbalance in GPL species has also been reported in other RBC disorders, including Chorea-Acanthocytosis and sickle cell disease [25–28]. In bone marrow (BM) failure disorders that are frequently associated with erythroid cytopenia, including myelodysplastic syndromes (MDS) and Diamond Blackfan anemia (DBA), metabolomic analyses have revealed abnormalities in erythroid cell lipid composition [29–31].

Erythropoiesis progresses through a hierarchical developmental pathway in which hematopoietic stem cells (HSC) form multipotent progenitor (MPP) that sequentially develop into common myeloid progenitor (CMP) and bipotent megakaryocyte-erythroid progenitor (MEP) [32–34]. The subsequent phase, early erythropoiesis (EE), involves the formation of erythroid progenitors burst-forming unit-erythroid (BFU-E) and colony-forming unit-erythroid (CFU-E) from MEP. Following this, terminal erythroid maturation (TED) involves passage through erythroblast stages, including proerythroblast (ProE), basophilic erythroblast (BasoE), polychromatic erythroblast (PolyE), and orthochromatic erythroblast (OrthoE) [35–43]. Finally, maturation is characterized by enucleation and considerable remodeling of the plasma membrane to achieve the characteristic biconcave RBC morphology [44,45]. Lipid composition changes include reduction in GPLs and enrichment of SLs, cholesterol, and PE during the late developmental stages [10,46,47]. Put together, there is strong evidence that membrane lipid composition is dynamically regulated during erythroid differentiation and its disruption is linked to defects in RBC homeostasis, however, molecular mechanisms underlying regulation of lipid profiles during normal erythropoiesis remain largely undefined.

In this study, we identify a novel role for erythropoietin (Epo) in regulating the lipid composition of developing erythroid cells. Epo is an established driver of the EE phase and essential for the survival and continued differentiation of BFU-E and CFU-E cells [35–42,48–51]. Using scRNA-seq of *ex vivo* cultures of BM CD34^+^ hematopoietic stem and progenitor cells (HSPCs) grown in the presence or absence of Epo, we delineated stage specific transcription dynamics associated with the BFU-E to CFU-E transition. In addition to the canonical Epo-responsive programs involved in heme and globin synthesis, iron homeostasis, and erythroid transcription factor regulation, gene set enrichment analysis (GSEA) revealed the involvement of lipid metabolic pathways in the BFU-E to CFU-E transition. Further, untargeted lipidomics revealed Epo-driven remodeling of cellular GPL species, indicating metabolic reprogramming. Integration of lipidomic and transcriptomic data uncovered Epo-mediated regulation of GPL biosynthetic enzymes, which was validated at the protein level using intracellular flow cytometry. Together, these findings demonstrate Epo’s critical role in lipid homeostasis, providing new insights into mechanism by which membrane lipid profiles are shaped in erythroid cells.

## Materials and Methods

### Single Cell RNA Sequencing and Data Analysis

Human BM-derived CD34^+^ cells, all from female donors between the ages 13 to 34 years (Ossium Health, San Francisco, CA, USA), were cocultured with mouse MS-5 stromal cells in medium with or without 4U/mL Epo (+/−Epo conditions) for seven days as described previously [52–55]. Libraries were prepared from cells from two donors using the Chromium Next GEM Single Cell 3’ Reagent Kit v3.1 (10x Genomics) according to the manufacturer’s instructions and sequenced using the Novogene platform (paired-end sequencing, 150 bp reads).

CellRanger v7.0.1 (10x Genomics) was used for the processing of raw FASTQ files, mapping reads to the human genome (GRCh38-2020-A), and generation of unique molecular identifier (UMI) counts. Seurat version 4.4.0 was used to import data for all four samples and to perform quality control, log normalization, and unsupervised clustering, and DoubletFinder was used for doublet removal. Seurat was also used to merge samples, filter out clusters, and perform DEG analysis. Clusters were manually annotated using established markers; see Supplemental Materials and Methods and Supplemental File 2 for details. ShinyCell and Monocle3 were used for data visualization and pseudotime analysis, respectively. GSEA was performed using the WEB-based Gene Set Analysis Toolkit (WebGestalt) [56]. The list of software packages used in this analysis is provided in Supplemental File 1.

### Lipid Extraction and Mass Spectrometry-Based Lipidome Analysis

Cells from BM CD34^+^ HSPC cultures (four different donors), grown with and without Epo, were harvested on day 7, and lipid metabolites were extracted using methanol (MeOH; LC/MS Grade) and tert-butyl methyl ether (MTBE). Lipidome analysis was performed by liquid chromatography and tandem mass spectrometry (LC-MS/MS) on a Vanquish UPLC-Exploris 240 Orbitrap MS. For each sample, the intensities of identified metabolites were divided by the total number of cells, and duplicate metabolites within samples were summed. Statistical analysis was performed in MetaboAnalyst 6.0, and prediction of lipid reactions and pathways was performed using the BioPAN tool within LIPID MAPS [57].

### Intracellular Immunophenotyping

BM HSPCs cultured in medium with and without Epo were collected, incubated first with the fixable viability stain Live-or-Dye™ 405/545 (Biotium) and then stained with an antibody cocktail including lineage (Lin) markers CD3, CD10, CD11b, CD19, and CD41a, as well as the progenitor marker CD123, all conjugated to PerCP-Cy5.5, CD34-APC-Cy7, CD71-APC, and CD105-BV421. Next, cells were fixed with 1% paraformaldehyde, permeabilized with ice-cold 70% EtOH, and then blocked. Finally, cells were stained with unlabeled rabbit-anti-human primary antibodies against intracellular proteins, followed by DyLight 488 goat-anti-rabbit secondary antibody, and analyzed by flow cytometry. The list of antibodies used for analysis is provided in Supplemental File 2.

### Statistical Analysis

Graphical representation plots and statistical analysis by student T-test were performed in GraphPad Prism (version 10.0.3; La Jolla, CA). Data is represented as mean ± standard error of the mean (SEM), and p-values <0.05 were considered significant.

## Results

### Development of Single Cell Transcriptome-Based Erythroid Differentiation Trajectory

To investigate transcriptional dynamics that drive stage-specific Epo-dependent transitions within EE populations, we performed scRNA-seq. BM CD34^+^ HSPCs from two healthy donors were cultured for seven days in medium with and without Epo as described previously [52–55]. To avoid perturbations caused by cell sorting, harvested cells were directly used for the preparation of scRNA-seq libraries and sequenced up to a depth of ∼30,000 reads per cell. After sequence mapping, quality control, and normalization, the transcriptomic datasets from all samples were merged in Seurat [58]. Unsupervised clustering of the dataset from 26,387 cells yielded an all-cell UMAP containing 15 clusters that were annotated based on expression of established stem, progenitor, and lineage markers (Figure 1A and S1; see Supplemental Materials and Methods and Supplemental File 3 for details). To focus on transcriptomic changes associated with erythroid differentiation, clusters containing granulocytes (cluster 4), monocytes (cluster 7), megakaryocytes (clusters 11 and 12), and GMPs (cluster 10), as well as the two island clusters, 13 and 14, that contained only 209 and 158 cells, respectively, were filtered out. Unsupervised reclustering of the remaining subset, consisting of 19,231 cells, generated an erythroid UMAP of eleven clusters (Figure 1B). Pseudotime analysis using Monocle3 provided the cluster progression from least to most mature as C0→C1→C9→C2→C3→C4→C10→C6→C8→C5→C7 (Figures 1C and 1D).

**Figure 1.**
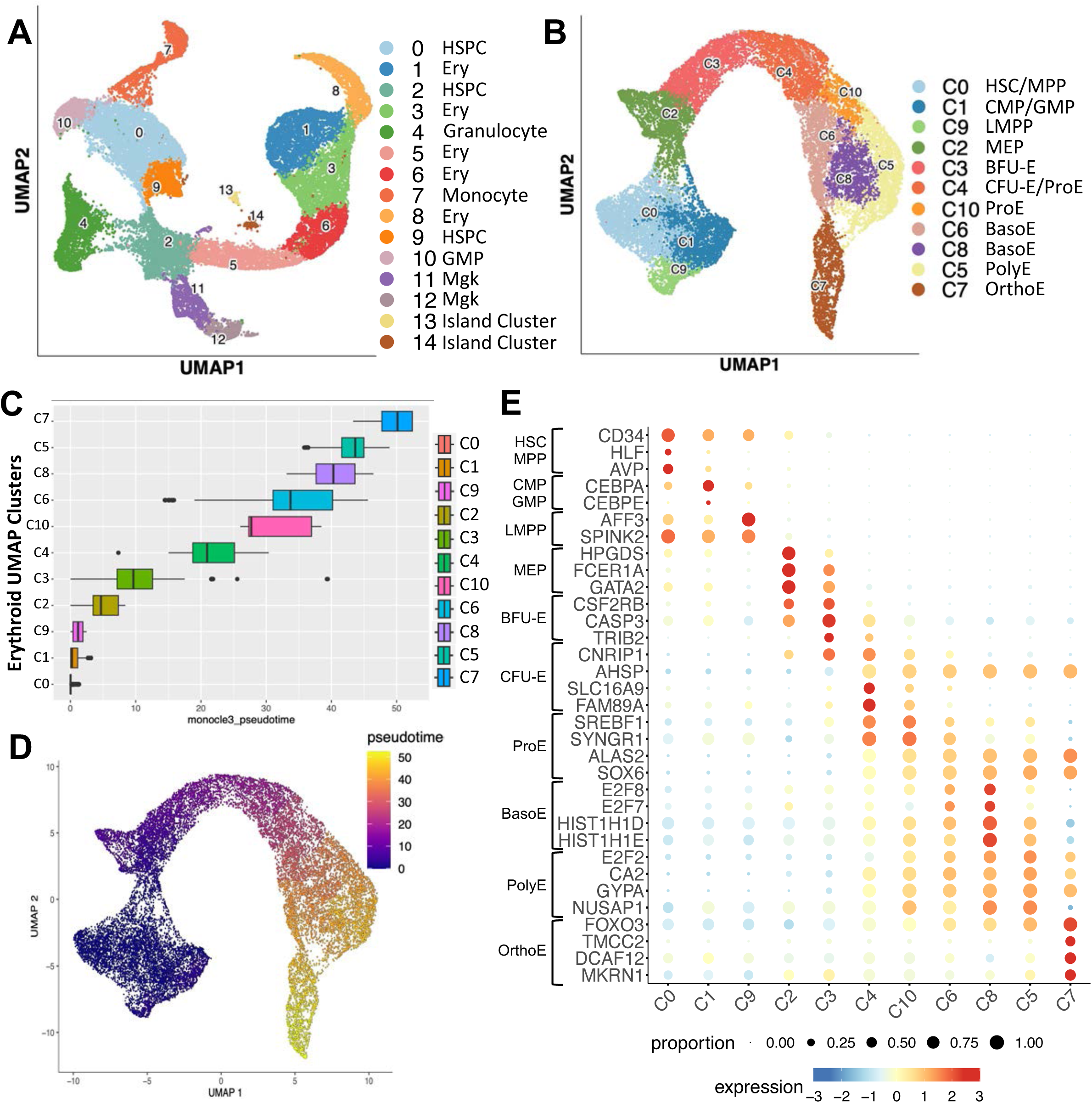
Single cell transcriptional profiling of cells from BM HSPC cultures. ScRNA-seq was performed on BM CD34^+^ HSPC cultures from two donors grown for 7 days in the presence and absence of Epo. **(A)** UMAP visualization of unsupervised clustering of all cells, identifying clusters specific to stem, progenitor, and differentiated lineage populations. **(B)** UMAP visualization of stem, progenitor, and erythroid populations. From the all cell UMAP, clusters containing granulocytes (4), monocytes (7), GMP (10), and megakaryocytes (11 and 12), and two island clusters (13 and 14) were filtered out to generate an erythroid UMAP. See Figure S1 for details cluster annotation. **(C)** Boxplot depicting the distribution of cells in psuedotime, grouped by cluster. **(D)** UMAP visualization of cells colored by position in pseudotime using Monocle3. **(E)** Bubble plot depicting expression of representative marker genes used for cluster annotation. Dot size represents the proportion of cells that express each marker, and the color represents scaled expression. Abbreviations: HSPC; hematopoietic stem and progenitor cell, Ery; erythroid, GMP; granulocyte monocyte progenitor, Mgk; megakaryocyte, HSC; hematopoietic stem cell, MPP; multipotent progenitor, CMP; common myeloid progenitor, LMPP; lympho-myeloid primed progenitor, MEP; megakaryocyte-erythroid progenitor, BFU-E; burst-forming unit erythroid, CFU-E; colony-forming unit erythroid, ProE; proerythroblast, BasoE; basophilic erythroblast, PolyE; polychromatic erythroblast, OrthoE; orthochromatic erythroblast.

We used the expression of representative marker genes, along with the pseudotime order, for manual annotation of the eleven clusters (Figures 1B-E, S2, and Supplemental File 3). Clusters C0, C1, and C9 were found to contain HSCs/MPPs, CMPs/GMPs, and lympho-myeloid primed progenitors (LMPPs), respectively, and C2 was found to be comprised of MEPs. The expression patterns of EE progenitor and erythroblast specific genes identified C3 as BFU-E, C4 as CFU-E and ProE, C10 as ProE, C6 and C8 as BasoE, C5 as PolyE, and C7 as OrthoE (Figure 1B).

### Delineation of the Epo-dependent Transition within Transcriptomic Clusters

Next, to uncover Epo-dependent transcriptional dynamics, we examined the erythroid UMAP to establish the transition beyond which clusters contained cells from +Epo cultures and not from −Epo cultures. All clusters were present in +Epo cultures from both donors at comparable percentages; however, clusters C4, C10, C6, C8, C5, and C7 were not observed in −Epo cultures (Figures 2A-B). This indicated a clear Epo-dependent step at the C3 to C4 transition, which also coincides with a rise in the expression of CD235a (Figure 2C). To assess transcriptional changes associated with this Epo-dependent transition, we first subsetted clusters C3 and C4 and then performed DEG analysis based on original culture condition (+/−Epo) (Supplemental File 4). Assessment of the top 400 DEGs (200 upregulated and 200 downregulated) revealed a switch in expression pattern between clusters prior to the onset of Epo-dependence (C0, C1, C9, C2, and C3) and those that follow (C4, C10, C6, C8, C5, and C7) (Figure 2D). The DEG list includes many previously reported Epo-dependent and erythropoiesis regulatory genes, such as the three erythroid master transcription factors (*GATA1*, *TAL1*, and *KLF1*), genes involved in heme and hemoglobin synthesis (*HBA2*, *HBA1*, *AHSP*, *HBG1*, *HBB*, etc.), and several others, such as *EPOR*, *ERFE*, *HES6*, and *JAK2* (Figure S3 and Supplemental File 4). Upregulation of these genes was observed in cells grown in the presence of Epo and in clusters that arise after the BFU-E (C3) to CFU-E (C4) transition. Additional details regarding the established erythropoiesis genes shown in Figure S3 are provided in the Supplemental Materials and Methods section.

**Figure 2.**
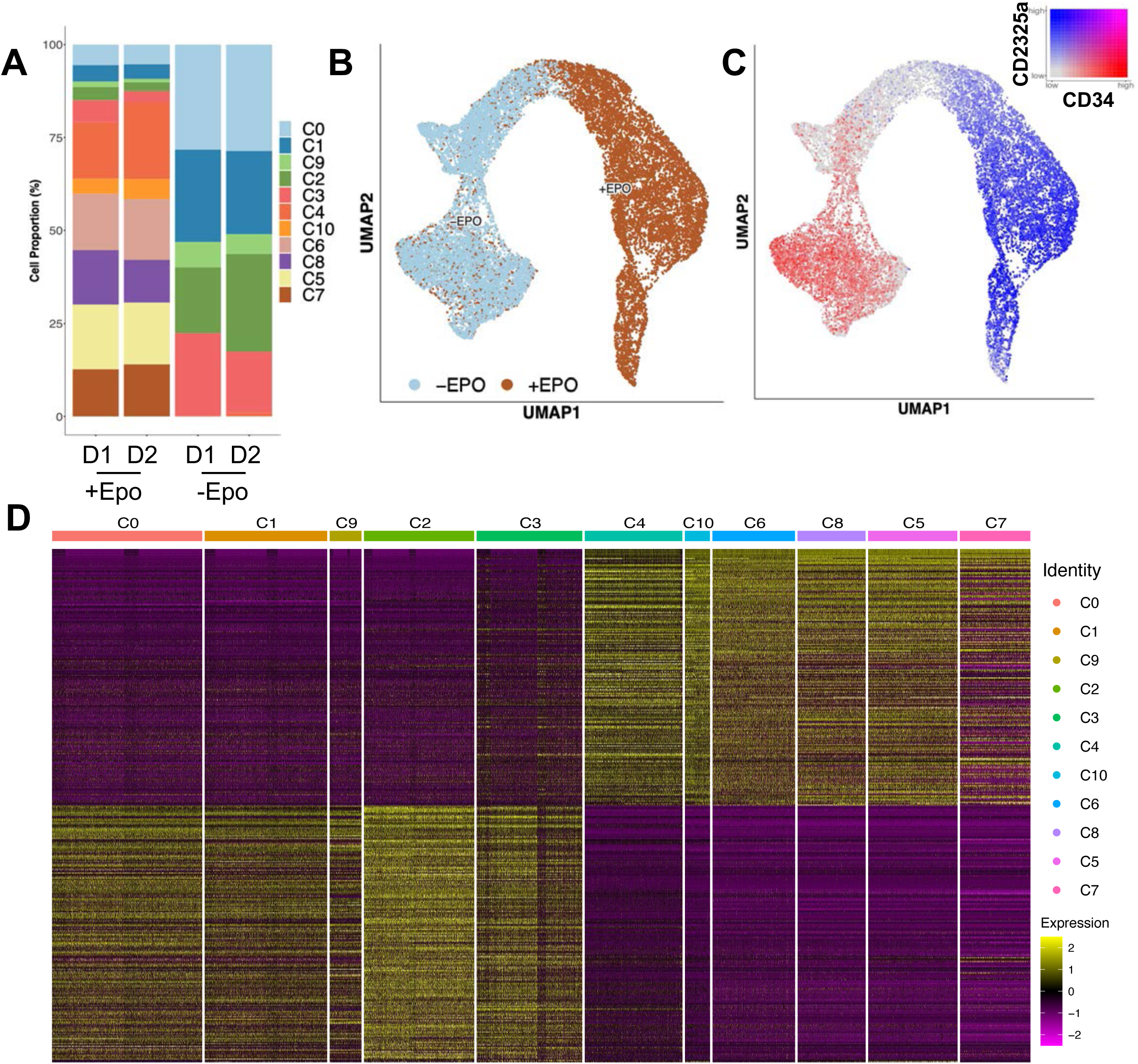
Epo-dependent transition during erythroid differentiation of human BM HSPCs. **(A)** Bar graph showing the proportion of cells in each cluster as a percent of total cells for all four samples. **(B)** UMAP visualization of clustered cells by original culture condition, +Epo or −Epo, highlighting an Epo-dependent transition at the threshold between clusters C3 and C4. **(C)** UMAP depicting coexpression of *CD34* and *GYPA* (CD235a). **(D)** Heatmap representation of expression of the top 200 up– and downregulated Epo-dependent DEGs (400 total), as observed in all clusters ordered in pseudotime. Columns represent clusters ordered according to pseudotime. Abbreviations: Epo; erythropoietin, D1; donor 1, D2; donor 2.

Previous transcriptional analyses in mice have found an Epo-associated upregulation of erythroid-related genes in MPP [59,60]. To examine if similar Epo-driven transcriptional changes occur in multi– and bipotent progenitors from *ex vivo* cultures of human cells, we subsetted HSPC and LMPP clusters, C0, C1, and C9 together, and the MEP cluster C2 separately. DEG analysis based on Epo treatment for both subsets revealed differential regulation of a number of genes (Supplemental File 5). Notably, seven of the top 10 upregulated genes were shared in both cohorts; these included globin-associated genes, *HBB*, *HBA2*, *HBA1*, *AHSP, HBG1*, and *HBM*, as well as *SLC25A37*, which is a mitochondrial iron transporter. These findings indicate that, consistent with previous reports [59,60], Epo elicits limited yet detectable transcriptional changes in hematopoietic progenitors.

### Resolution of Early Erythroid Heterogeniety Based on CD71 and CD105 Expression

In a previous study, we found an Epo-induced upregulation of CD71 (transferrin receptor) and CD105 (endoglin) in EE cells and reported that, based on CD34, CD71, and CD105, heterogenous EE cells could be immunophenotypically resolved into five CD235a^−^ subpopulations [55]. These include two CD34^+^ BFU-E subtypes—early BFU-E (earlyB; CD34^+^CD71^lo^CD105^lo^) and late BFU-E (lateB; CD34^+^CD71^hi^CD105^lo^)—and three CD34^−^ CFU-E populations—early CFU-E (earlyC; CD34^−^CD71^lo^CD105^lo^), mid CFU-E (midC; CD34^−^CD71^hi^CD105^lo^), and late CFU-E (lateC; CD34^−^CD71^hi^CD105^hi^). As such, the lateC population was unique in that it exhibited high expression of both CD71 and CD105 and was the only EE subtype that required Epo for its formation. Since the Epo-dependent transition in the scRNA-seq dataset was between clusters C3 and C4, it is plausible that lateCs are located in C4 and the remaining BFU-E (earlyB and lateB) and CFU-E (earlyC and midC) subtypes are in C3.

To examine if heterogeneity in EE cells could be resolved transcriptionally, we subsetted clusters C2 (MEP), C3 (BFU-E), C4 (CFU-E/ProE), and C10 (ProE) from the erythroid UMAP and reclustered them, yielding an EE UMAP of seven clusters (Figure 3A). In this UMAP, an Epo-dependent transition was observed between E3 and E2 (Figure 3B). Pseudotime analysis provided the progression order from least to most mature cluster as E0→E1→E6→E3→E2→E5→E4 (Figure 3C-D). Expression of the MEP marker *HPGDS* suggests that MEPs within cluster C2 of the erythroid UMAP (Figure 1B) segregated into two clusters in the EE UMAP: E0 and E6 (Figure 3A and S4A-B). While E0 likely represents true MEPs, E6 appears to be comprised of megakaryocyte-primed MEPs because it exhibits high expression of megakaryocyte markers, *FL1* and *CD9* (Figure 3A and S4C-D). The pattern of BFU-E (*TRIB2*) and CFU-E (*FAM89A*) markers suggests that clusters C3 and C4 also segregated into two clusters each: C3 into E1 and E3, and C4 into E2 and E5 (Figure S4E-F). The rise in expression of CD235a and other ProE markers *SOX6* and *ALAS2* indicates that E5 and E4 are both ProE clusters (Figure 3A and S4G-I). Thus, this analysis segregates CFU-E from ProE clusters. Further, Epo-dependence of E2 formation indicates that the CD71^hi^CD105^hi^ lateCs are located in E2, and other EE populations lie within E1 and E3.

**Figure 3.**
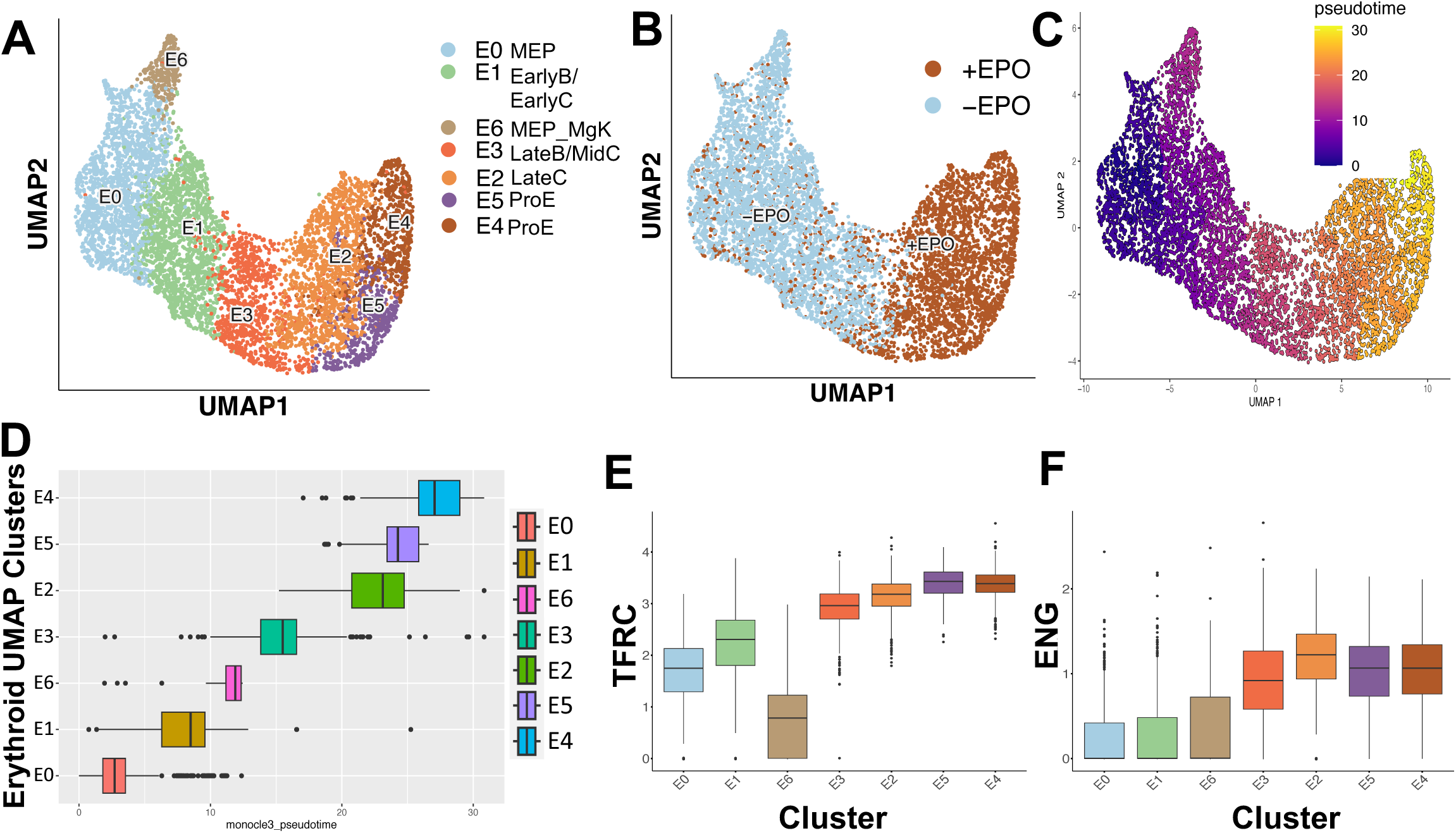
Transcriptional resolution of early erythroid populations. From the erythroid UMAP, clusters C2 (MEPs), C3 (BFU-E), C4(CFU-E/ProE), and C10 (ProE) were subset and re-clustered to further resolve EE subtypes. **(A)** UMAP visualization and annotation of EE clusters. **(B)** UMAP visualization of clustered cells by original culture condition, +Epo or −Epo, highlighting an Epo-dependent transition at the threshold between clusters E3 and E2. **(C)** UMAP visualization depicting the progression of cells in pseudotime using Monocle3. **(D)** Boxplot depicting the distribution of clusters in psuedotime. Boxplots depicting gene expression of **(E)** *TFRC* (CD71) and **(F)** *ENG* (CD105) in clusters ordered in pseudotime. Abbreviations: MEP; megakaryocyte-erythroid progenitor, MgK; megakaryocyte, EarlyB; early BFU-E, LateB; late BFU-E, EarlyC; early CFU-E, MidC, mid CFU-E, LateC; late CFU-E, ProE; proerythroblast Epo; erythropoietin.

Assessment of *TFRC* (CD71) revealed its high expression across all clusters other than the megakaryocyte-primed MEP cluster E6; however, there was a noticeable rise in its expression levels between E1 and E3 (Figure 3E). This suggests that lateB and midC, which immunophenotypically exhibited high expression of CD71, are likely located in E3, whereas CD71^lo^ earlyBs and earlyCs are in E1. The highest expression of *ENG* (CD105) in the Epo-dependent E2 further supports the location of lateCs in this cluster (Figure 3F). Thus, similar to the immunophenotyping analysis, the current mRNA quantification underscores the heterogeneity of developing EE cells and discerns the most probable distribution of the five EE populations based on CD71 and CD105.

### Identification of Epo-dependent Molecular Processes During Erythroid Transitions

Next, we used Seurat to analyze DEGs in all eleven clusters of the erythroid UMAP to identify additional cluster-specific genes and Kyoto Encyclopedia of Genes and Genomes (KEGG) pathways underlying the successive transitions (Figure 1B). Expression of the top 20 DEGs within C0, C1, and C9, containing HSC/MPP, CMP/GMP, and LMPP, respectively, showed overlapping profiles (Figure 4A and Supplemental File 6). The MEP cluster C2 appears to be distinct from other HSPCs but overlaps with the BFU-E cluster C3. Similarly, the CFU-E cluster C4 is distinct from BFU-Es but exhibits profile overlap with ProE containing C10. Among clusters spanning erythroblast stages, the transcriptional profile of C10 exhibits similarity with C4 and C6, and that of C8 exhibits similarity with C6 and C5. These patterns indicate a gradual activation and repression of stage-specific genes, suggesting an erythroblast continuum. OrthoEs, however, form a distinct cluster (C7) and exhibit a unique expression profile.

**Figure 4.**
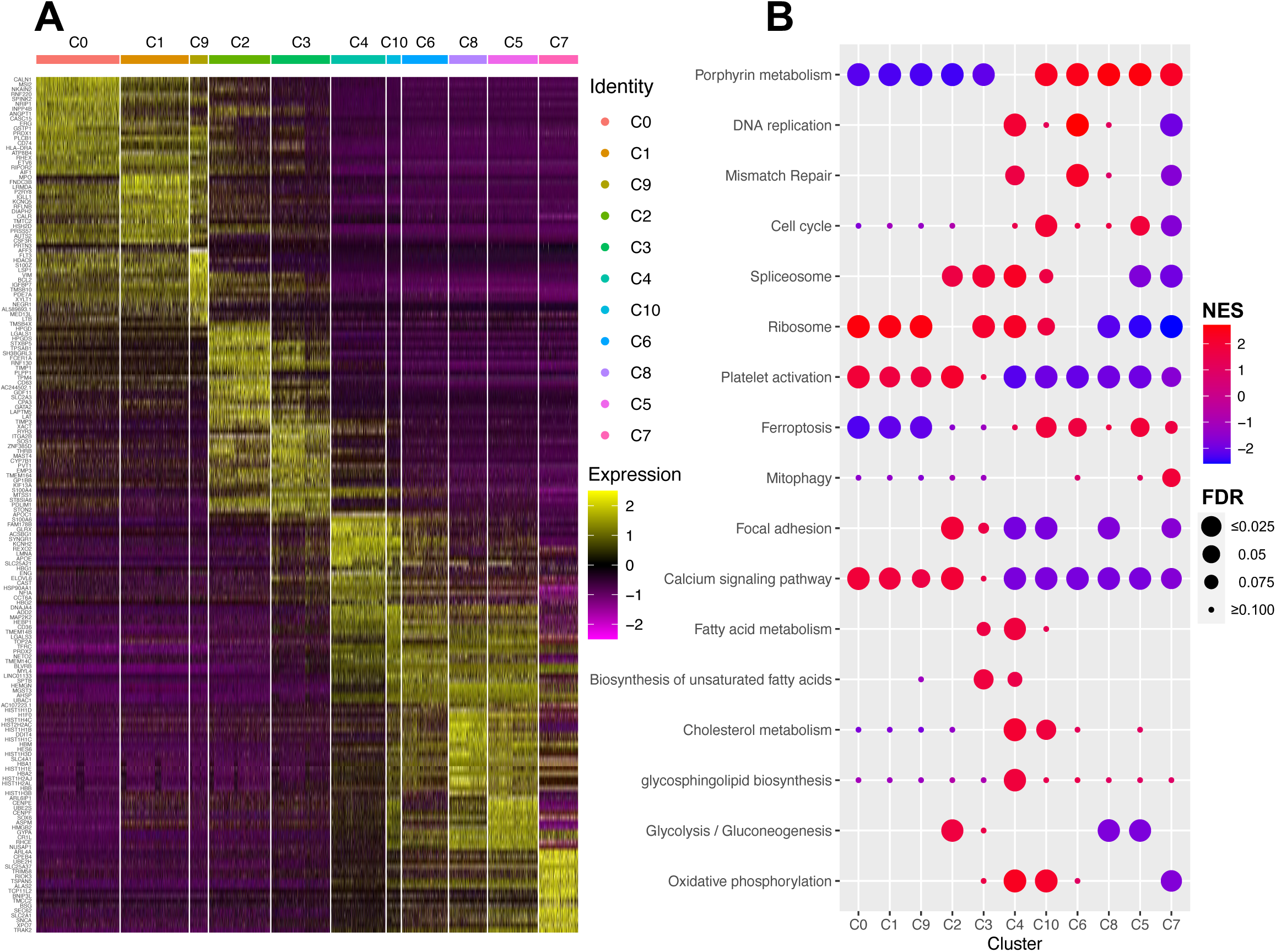
Epo-dependent differentially expressed genes and gene ontology analysis. **(A)** Heatmap depicting expression trends of the top 20 differentially upregulated genes in each cluster. Columns are ordered according to pseudotime showing progression from immature to mature clusters. **(B)** Bubble plot of terms found to be enriched in each cluster by GSEA. The dot color represents the normalized enrichment score (NES), and the size indicates the false discovery rate (FDR).

To identify cellular pathways that drive Epo-dependent transition, we performed GSEA [56]. For this, the entire DEG list for clusters C3 and C4 was used as an input into GSEA, and predominant KEGG pathways were assessed across all clusters (Figure 4B). This analysis confirmed the involvement of several well-established erythroid differentiation pathways [10,43,61–63]. Top amongst these is protoporphyrin metabolism, which is repressed in C3 but significantly upregulated from C10 onwards. Upregulation of DNA replication, mismatch repair, and cell cycle, and downregulation of focal adhesion and calcium signaling are in alignment with the known expansion of erythroblasts during TED. Similarly, upregulation of RNA processing complexes—spliceosome and ribosome—indicates the demands of proliferating progenitors. All these processes, except protoporphyrin metabolism, decline later in erythroblasts. Other expected alterations include a downregulation of platelet activation and a slight upregulation of ferroptosis and mitophagy in erythroblast clusters.

Very interestingly, we found transient upregulation of lipid metabolism pathways in EE cells. Biosynthesis of unsaturated fatty acids and fatty acid metabolism were enriched in C3 and C4, cholesterol metabolism was up in C4 and C10, and glycosphingolipid (GSL) biosynthesis was up in C4 of the erythroid UMAP (Figure 4B). Other interesting metabolic events include a transient reduction in glucose metabolism and oxidative phosphorylation in late erythroblast-specific clusters (C8, C5, and C7), which have recently been reported to occur in mouse and human erythroblasts in other studies [10,62,64].

To further assess association with lipid metabolism pathways, we performed GSEA analysis on DEGs from the EE UMAP and observed an upward change not only in the four terms identified in Figure 4A, but also in glycosylphosphatidylinositol (GPI)-anchor biosynthesis, fatty acid elongation, and fatty acid biosynthesis at the Epo-dependent transition between E3 and E2 (Figure S5A). Together, the scRNA-seq and GSEA analyses revealed transcriptional alterations across all erythroid differentiation stages and identified the potential involvement of lipid metabolism pathways in Epo-dependent transitions.

### Epo Mediates Alterations in Lipid Profiles of Differentiating Cells

To examine the nature of Epo-induced alterations in lipid profiles of differentiating cells, we performed untargeted lipidomics. BM CD34^+^ HSPCs from four healthy donors were cultured in media with and without Epo. To avoid artifacts due to FACS, lipids were extracted from all culture cells on day 7 and then analyzed by LC-MS/MS. This analysis identified 1789 lipid species from four lipid classes, including fatty acids, glycerolipids, SL, and GPL (Supplemental File 7). Notably, cholesterol and cholestrylesters were not detected. In addition to ester linkages, ether-linked (alkyl and alkenyl) fatty acids were also detected in all lipid classes detected. Upon unsupervised clustering based on lipid metabolite levels per cell, the samples clustered by treatment, showing similar Epo-induced lipid profiles (Figure 5A). Principal component analysis (PCA) also showed clustering of samples by treatment and indicated close clustering in the absence of Epo but some variation in the response of the biological replicates to Epo (Figure 5B). Volcano plot analysis revealed Epo-induced alterations mainly in GPL, which comprised 74% of the significantly altered species (Figure 5C-D and Supplemental File 7). Within GPLs, the majority of PC, PE, and phosphatidylinositol (PI) species were reduced in −Epo compared to +Epo condition (Figure 5C-D). Lysophospholipids (LPLs), on the other hand, were increased in the absence of Epo. The majority of the altered lipid species (∼85%) contained unsaturated fatty acids, underscoring the significance of unsaturated fatty acid biosynthesis as an Epo-dependent molecular process (Figure 4B and 5C).

**Figure 5.**
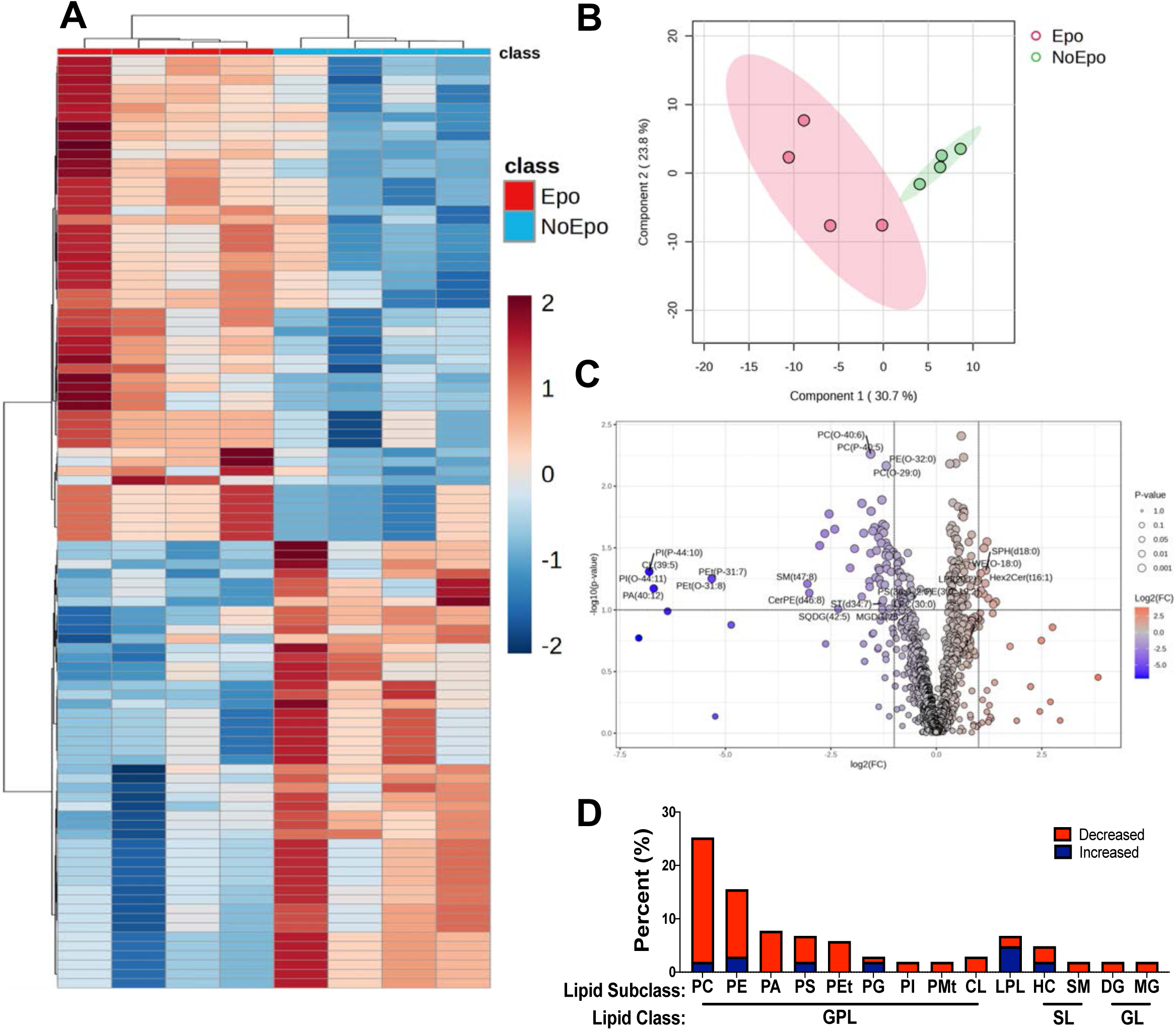
Untargeted analysis of erythropoietin mediated alterations in lipid profiles. Untargeted lipidomic analysis was performed on whole cell suspensions from BM CD34^+^ HSPCs from four donors cultured with (Epo) and without (NoEpo) Epo for seven days. **(A)** Heatmap depicting top 100 differentially regulated lipid metabolites. **(B)** Principal component analysis showing distribution of samples. Both (A) and (B) show unsupervised clustering of samples by treatment. **(C)** Volcano plot of differentially regulated lipid metabolites, comparing NoEpo vs Epo conditions, from which the list of significantly altered species was derived using a fold change (FC) > 2 or < 0.5 (Log2(FC) > |1|) (Supplemental File 7, “Significant Lipid Metabolites”). **(D)** Bar graph depiction of significant differentially regulated lipid subclasses, shown as a percentage of all significantly altered lipid species. Decreased and increased describe changes in levels observed in the absence of Epo (No Epo). Abbreviations: PC; phosphatidylcholine, PE; phosphatidylethanolamine, PA; phosphatidic acid, PS; phosphatidylserine, PEt; phosphatidylethanol, PG; phosphatidylgylcerol, PI; phosphatidylinositol, PMt; phosphatidylmethanol, CL; cardiolipin, LPL; lysophospholipid, HC; hexosylceramide, SM; sphingomyelin, DG; diglyceride, MG; monoglyceride, PL; phospholipid, SL; sphingolipid, GL; glycerolipid.

To gain an insight into lipid pathways and enzymes that may be associated with the Epo-induced alterations in lipid species, we used the BioPAN tool within LIPID MAPS [57]. Analysis of the lipidomic dataset predicted the highest activity score for the formation of PC from diacylglycerol (DG) and the lowest score for the conversion of PE to PC. Intermediate scores were predicted for the conversion of PC to PS and PS to PE and for the formation of PE from DG (Figure 6A). This analysis also predicted involvement of several GPL metabolism genes in either stimulation or suppression of specific reactions (Supplemental File 7). To examine correlations between BioPAN predictions and the scRNA-seq dataset, we assessed the expression of GPL metabolism genes across all clusters of the erythroid UMAP. This revealed differential Epo-mediated regulation of several genes; in response to Epo, *PTDSS1* and *2*, *MBOAT1*, *PEMT*, and *CEPT1* were downregulated, whereas *LPCAT3*, *CHPT1*, and *MBOAT2* were upregulated, and *PISD* was unaffected (Figure 6B). Importantly, these changes in expression also coincided with the BFU-E (C3) to CFU-E (C4) transition (Figure 6C). Furthermore, examination of the EE UMAP revealed similar alterations in these GPL biosynthesis genes, showing Epo-dependent changes in GPL metabolism genes at the level of both sample and cluster (Figure S5B-C).

**Figure 6.**
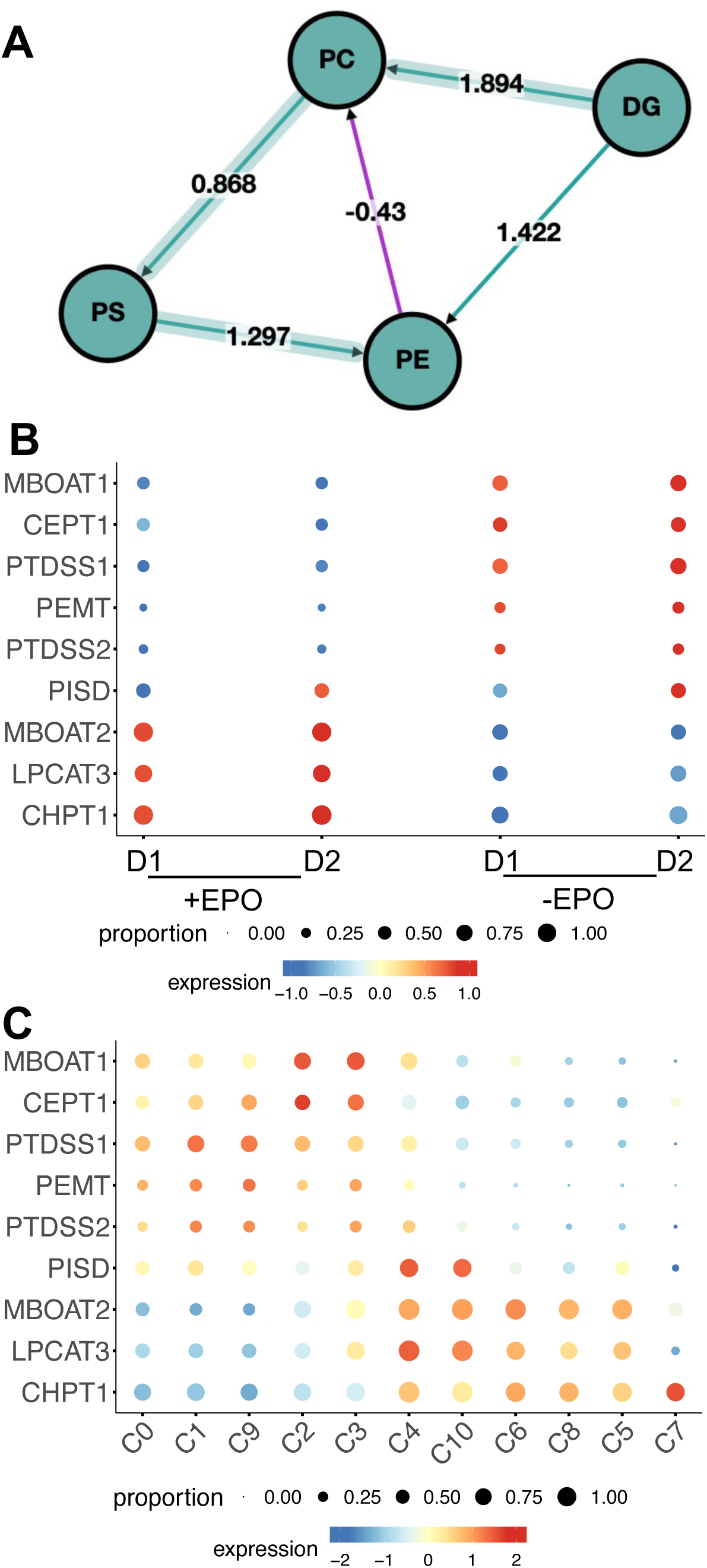
Transcriptional profiles of Epo-dependent glycerophospholipid metabolism genes. **(A)** Schematic representing BioPAN analysis of the stimulated or suppressed GPL metabolic reactions. Green nodes represent active lipids, and green shaded arrows represent active pathways resultant of comparing Epo vs No Epo. P-value < 0.05 equivalent to Z-score >|1.645|; represents significantly altered reaction between the Epo and NoEpo groups; Green arrow: positive Z-score; purple arrow: negative Z-score. **(B)** Bubble plot depicting relative expression of GPL metabolism genes that were predicted to be altered in an Epo-dependent manner in BioPAN analysis. **(C)** Cluster specific expression of Epo-dependent GPL metabolism genes. Dot color represents normalized expression, and size indicates the proportion of cells that express the gene within a cluster. Abbreviations: PC; phosphatidylcholine, PE; phosphatidylethanolamine, PS; phosphatidylserine, DG; diacylglycerol, Epo; erythropoietin, D1; donor 1, D2; donor 2, MBOAT(1 or 2); membrane bound glycerophospholipid O-acyltransferase (1 or 2), CEPT1; choline/ethanolamine phosphotransferase 1, PTDSS(1 or 2); phosphatidylserine synthase (1 or 2), PEMT; phosphatidylethanolamine N-methyltransferase, PISD; phosphatidylserine decarboxylase, LPCAT3; lysophosphatidylcholine acyltransferase 3, CHPT1; Choline phosphotransferase 1.

As observed above, the Epo-dependent transition was associated with a transient upregulation in GO terms associated with lipid metabolism. While we see significant changes in the expression of GPL metabolism gene when analyzed by treatment (+/−Epo) (Figure 7A), GPL metabolism did not appear as a significant term in GSEA. Untargeted lipidomic analysis of HSPC cultures, however, identified GLPs and LPLs as the most down– and upregulated lipid species, respectively, thereby implicating GPL metabolism in Epo-mediated erythroid differentiation (Figure 5C-D). To validate differential expression of GPL metabolism genes at the protein level, we performed intracellular staining and flow cytometry. Cells from BM-derived CD34^+^ HSPC cultures, grown in the presence and absence of Epo for seven days, were harvested, stained with anti-CD34, –CD71, –CD105, –CD123, and Lin antibodies, fixed, permeabilized, and then incubated with unconjugated rabbit polyclonal antibodies to GPL metabolism enzymes followed by DyLight488-conjugated anti-rabbit secondary IgG. An unconjugated rabbit polyclonal anti-GFP antibody was used as a negative control and to determine gates for positive cells. Additionally, SF3A1, a splicing factor whose expression was unaltered during EE, was used as a positive control (Figure 7A). For analysis, LiveLin^−^CD123^−^CD34^−^CD71^+^CD105^+^ cells were selected, and changes in percent positive cells and mean fluorescence intensity (MFI) for DyLight 488 were determined (Figure 7B-D). In agreement with scRNA-seq results, fold change in positive cells and MFI for LPCAT3, CHPT1, and MBOAT2 were significantly reduced for cultures grown in the absence of Epo (Figure 7). CEPT1 and PEMT, which were predicted to have higher expression in the absence of Epo, exhibited some increase in −Epo cultures; however, only PEMT was significantly altered (Figure 7). Together, these results suggest that Epo may be reprogramming GPL metabolism in differentiating erythroid cells within the EE phase.

**Figure 7.**
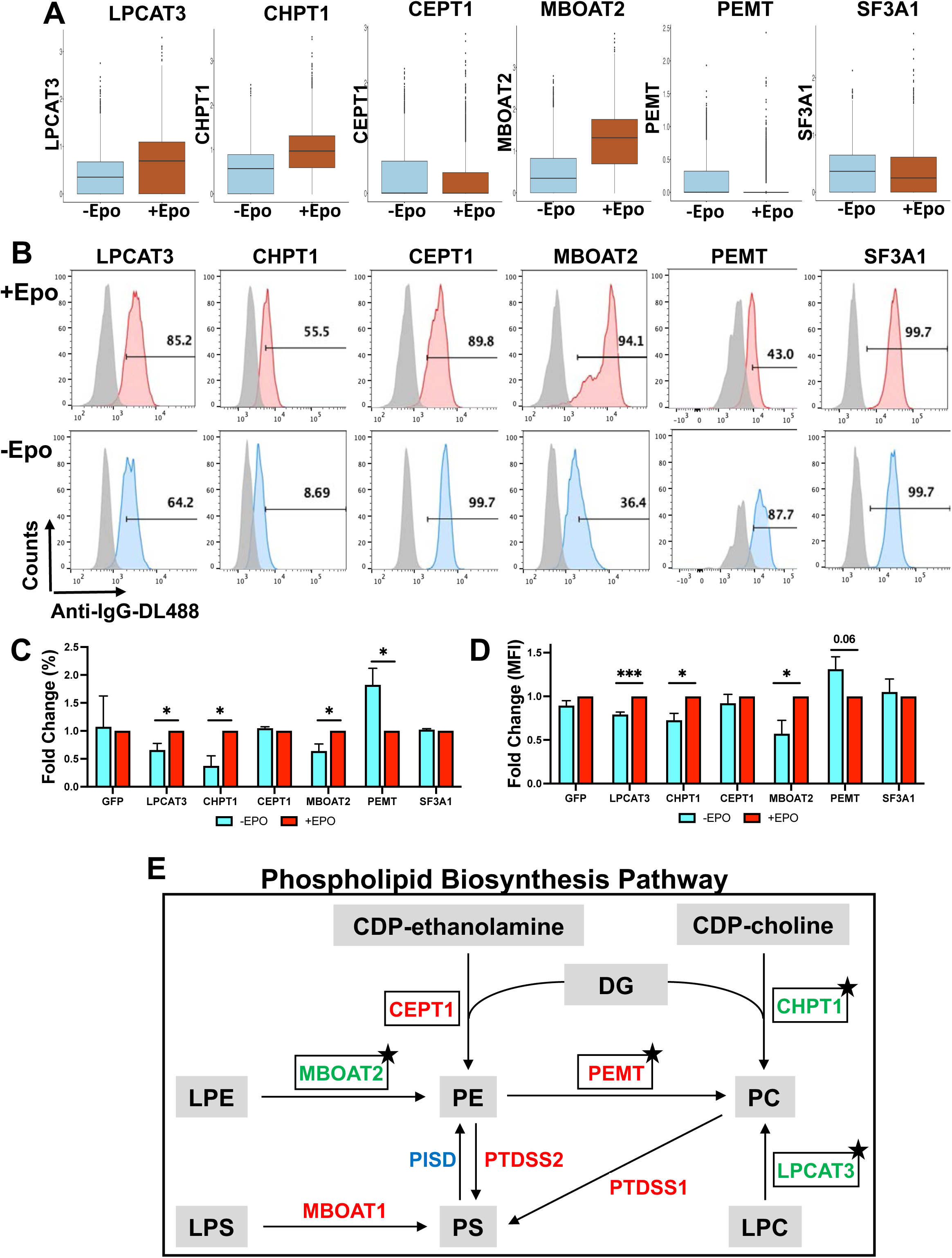
Validation of Epo-dependent changes in glycerophospholipid metabolism enzymes. BM-derived CD34^+^HSPCs were cultured for seven days in the presence (+Epo) and absence (−Epo) of Epo. Whole cell suspensions were collected, probed for surface and intracellular antibodies and analyzed by flow cytometry (see Materials and Methods). **(A)** Boxplots depicting expression pattern of transcripts from GPL metabolism genes in the scRNA-seq dataset. **(B)**Representative histograms showing expression of GFP (grey) and target antibody in cells from +Epo (top) and −Epo (bottom) cultures. **(C)** Relative fold change in the percentage of cells stained with antibodies against GFP, GPL metabolism enzymes (LPCAT3, CHPT1, CEPT1, MBOAT2, and PEMT), and SF3A1 expression (percent). **(D)** Relative fold change in MFI of cells stained with antibodies against protein or enzyme. Fold changes for both (C) and (D) were calculated in cells grown without Epo relative to with Epo. P-values ≤ 0.05 were considered significant (*p < 0.05; n = 3 or 4). **(E)** Simplified schematic of steps in GPL metabolism with the corresponding enzymes. Enzymes colored in red were found to be downregulated in +Epo cultures in our scRNA-seq dataset, while enzymes colored in green were upregulated, and PISD (blue) was unchanged. Squares indicate enzymes that were validated at the protein level and stars indicate the ones found to be significantly changed in cells cultured with Epo by intracellular flow cytometry. These results suggest Epo-mediated reprograming of GPL metabolism during erythropoiesis, particularly in the EE compartment. Abbreviations: Epo; erythropoietin, DL488; DyLight488; MFI; mean fluorescence intensity, CDP; cytidine di phosphate, PC; phosphatidylcholine, PE; phosphatidylethanolamine, LPC; lysophosphatidylcholine; PE; phosphatidylethanolamine, PS; phosphatidylserine, DG; diacylglycerol, LPE; lysophosphatidylethanolamine, LPS; lysophosphatidylserine, CHPT1; Choline phosphotransferase 1, LPCAT3; lysophosphatidylcholine acyltransferase 3, PEMT; phosphatidylethanolamine N-methyltransferase, CEPT1; choline/ethanolamine phosphotransferase 1, MBOAT; membrane bound glycerophospholipid O-acyltransferase, PISD; phosphatidylserine decarboxylase, PTDSS; phosphatidylserine synthase, SF3A1; splicing factor 3A subunit 1.

The major cellular pathway for *de novo* synthesis of PC is the cytidine diphosphate (CDP)-choline pathway, also known as the Kennedy pathway (Figure 7E). Choline phosphotransferase 1 (CHPT1), the final enzyme in this pathway, forms PC from CDP-choline and DG [65]. In addition, PC is also formed by the conversion of LPC to PC, a reaction catalyzed by lysophosphatidylcholine acyltransferase 3 (LPCAT3) through the Lands cycle, and by methylation of PE by phosphatidylethanolamine N-methyltransferase (PEMT). Similarly, PE can be formed from CDP-ethanolamine through the Kennedy pathway, catalyzed by choline/ethanolamine phosphotransferase 1 (CEPT1), and by the conversion of lysophosphatidyl ethanolamine (LPE) to PE, catalyzed by membrane-bound glycerophospholipid O-acyltransferase 2 (MBOAT2). Notably, CEPT1 favors PE formation but can also catalyze the formation of PC [66]. Our combinatorial omics analysis and follow-up validation by flow cytometry indicate a significant upregulation of CHPT1, LPCAT3, and MBOAT2 and downregulation of PEMT by Epo. This suggests that Epo-mediated reprogramming of GPL metabolism promotes the formation of PC and PE via distinct routes, enhancing *de novo* formation and acylation of LPC to PC but inhibiting the conversion of PE to PC (Figure 7E). Formation of PE by the conversion of LPE is also increased in the presence of Epo.

## Discussion

Anemia affects ∼25-30% of the global population and is a common symptom of many genetically inherited and acquired illnesses, underscoring the need for a comprehensive knowledge of erythropoiesis regulation [67,68]. Advanced technologies are facilitating a more nuanced understanding of processes driving erythroid commitment and maturation. While scRNA-seq analyses largely support the hierarchical framework of erythroid differentiation [69–72], they reveal alternative trajectories for progenitor development, including early erythroid priming from the most primitive HSCs and segregation of erythroid fate with other lineages [60,73–81]. Other studies have resolved erythroid heterogeneity and identified stage-specific genomic and protein markers [61,69,72,75,80,82–86]. Single cell analysis of *ex vivo* cultured human BM cells have unraveled how imbalance of heme and globin synthesis in CFU-E can induce a “death pathway”, characterized by upregulation of apoptosis, ferroptosis, and stress response genes [62]. Adopting a similar approach, we used scRNA-seq to analyze *ex vivo* cultures of human BM CD34^+^ HSPCs that were either supplemented with or lacking Epo. In addition to observing Epo-driven expression of several established erythroid pathways, we gained insight into the role of Epo in regulation of GPL metabolism. Furthermore, this analysis revealed Epo-associated alterations in erythroid genes even in stem and progenitor cells and recapitulated heterogeneity within EE progenitor cells.

Our single cell transcriptomic analysis uncovered a previously undescribed and vital role of Epo in transcriptional regulation of lipid metabolism in erythroid cells. Complementary lipidomics and protein validation indicate Epo-mediated reprogramming of GPL biosynthesis, leading to increased production of PC by the Kennedy pathway and increased acylation of LPC by Lands cycle, but decreased conversion of PE to PC (Figure 7E) [87]. We propose that this reprogramming is required to achieve the appropriate GPL composition needed to support the BFU-E to CFU-E transition within EE and/or early steps of TED. This idea is supported by studies that demonstrate link between abnormal lipid composition and defective erythroid differentiation in BM failure diseases, including MDS and DBA [29–31]. Others studies have reported decrease in GPL species abundance during reticulocyte maturation and shown that the breakdown of phosphocholine, formed during the degradation of PC to choline, is critical for erythroid differentiation in both mice and humans [46,88]. Thus, it is likely that Epo-mediated accumulation of PC in the membranes of EE progenitors is required to support downstream developmental transitions.

Due to lack of intracellular organelles, the capacity of mature RBCs to synthesize or breakdown major metabolites, including amino acids, nucleotides, and lipids is limited, however, during erythropoiesis many are critical for erythroid commitment and stage specific development [89–91]. While significant progress has been made in the understanding and targeting of the metabolic pathways that support ATP production, iron homeostasis, and heme biosynthesis, much remains to be done regarding regulation of membrane lipid profiles in erythroid cell development and maintenance. As such, further analysis is needed to determine what Epo-induced signaling pathways are involved in reprograming of the GPL biosynthesis pathway and what are the potential effects of alterations in GPL metabolism genes, identified in this study, on the development of BFU-Es and CFU-Es and, by extension, overall erythroid output. Furthermore, several studies, including ours, have implicated regulation of cholesterol, GSL, and fatty acid metabolism in the development and normal physiological functions of erythroid cells [12,92–95]. Thus, investigations are also needed to better understand the roles of these other lipid pathway genes in erythropoiesis. Insight into mechanisms of lipid metabolism regulation during erythroid development can potentially lead to targets that can be modulated to stimulate RBC production.

## Data Availability

The data underlying this article are available in Gene Expression Omnibus at https://www.ncbi.nlm.nih.gov/geo/, and can be accessed with accession number GSE278794.

## Acknowledgments

The authors would like to acknowledge funding support to SS from the National Institutes of General Medical Sciences (R01GM127464) and National Cancer Institute (P30CA023074) of the National Institutes of Health, the Valley Research Partnership Program (VRP P1-4009 and VRP77), and the Arizona Biomedical Research Center (ABRC: RFGA2022-010-30), to SQ from National Institute of Mental Health (R01MH128192) and National Eye Institute (NEI; R01EY035138), and to PS from NEI (R01EY025536). The content is solely the responsibility of the authors and does not necessarily represent the official views of the National Institute of Health. N.S. and S.S. would like to acknowledge the support provided by Mrinalini Kala, PhD Director of the Flow Cytometry Core.

## Figure Legends

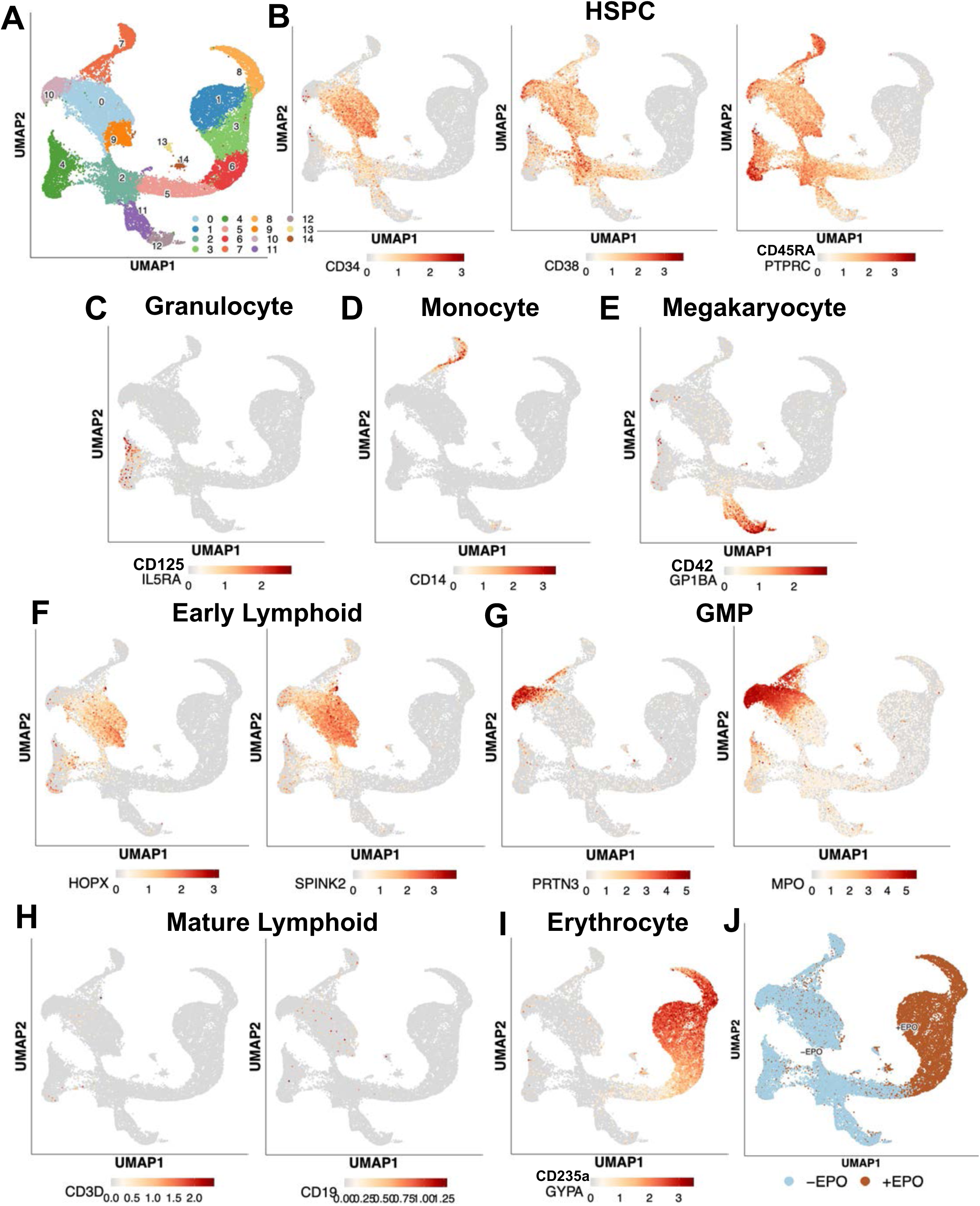
Figure S1.

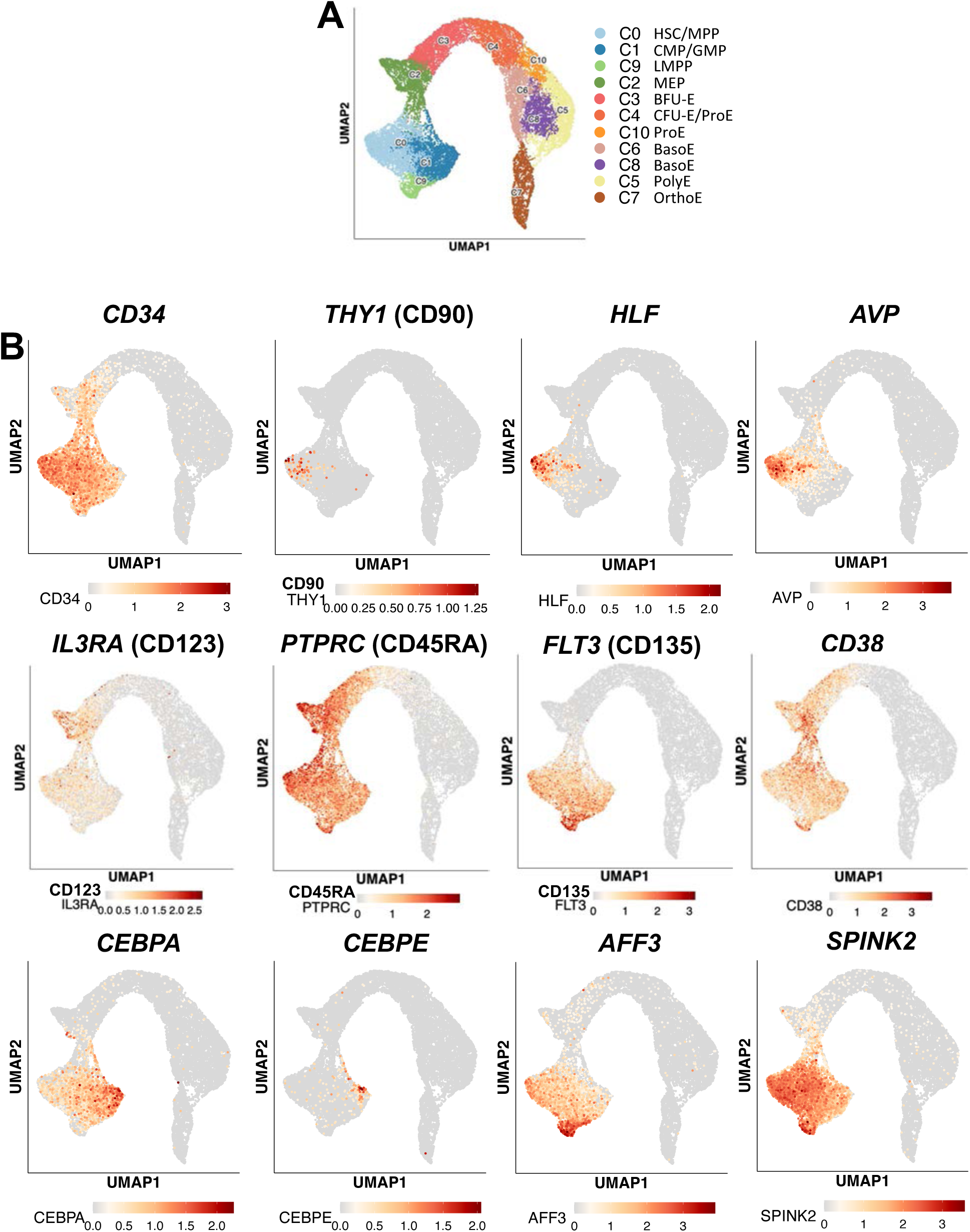
Figure S2.

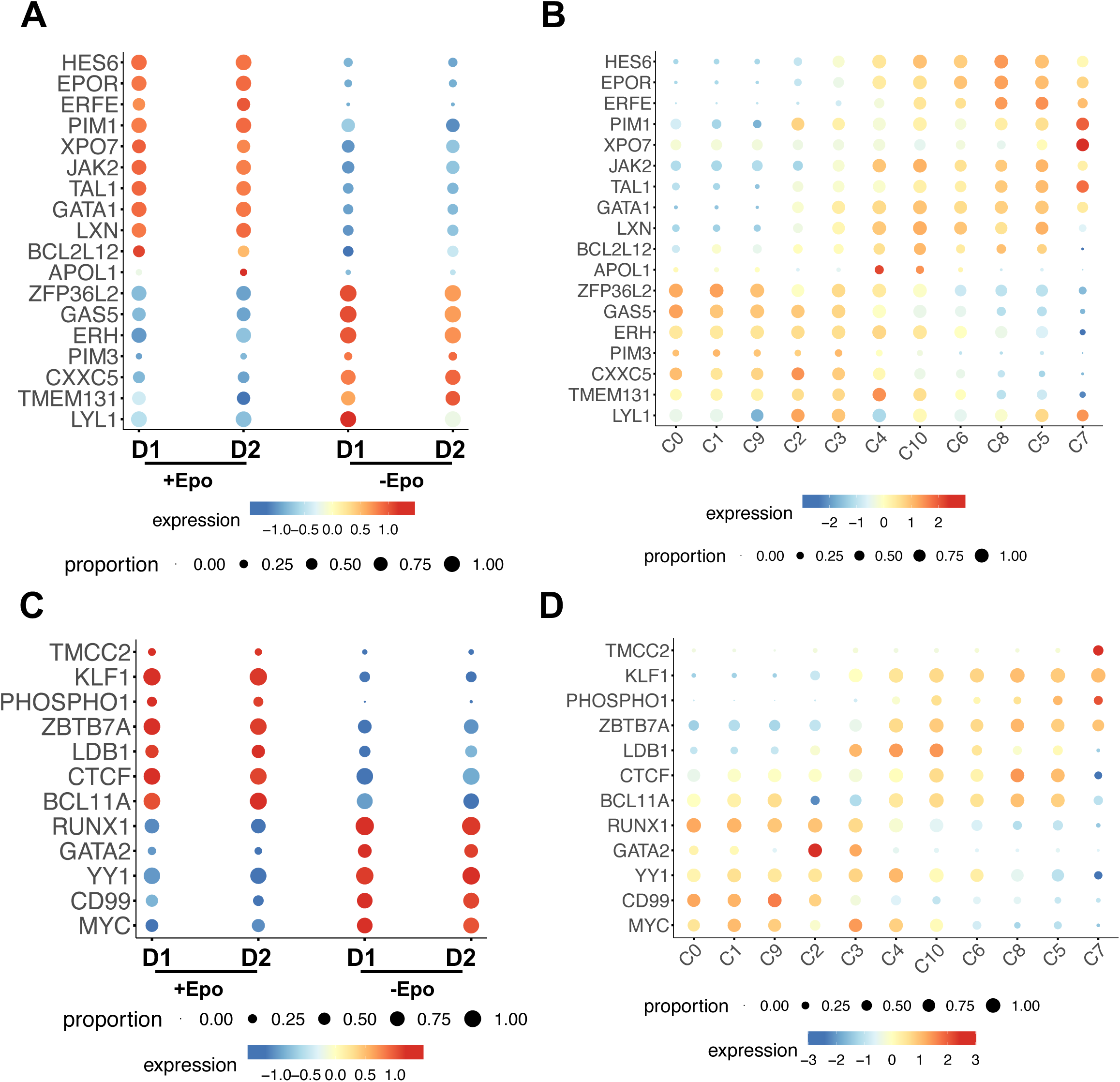
Figure S3.

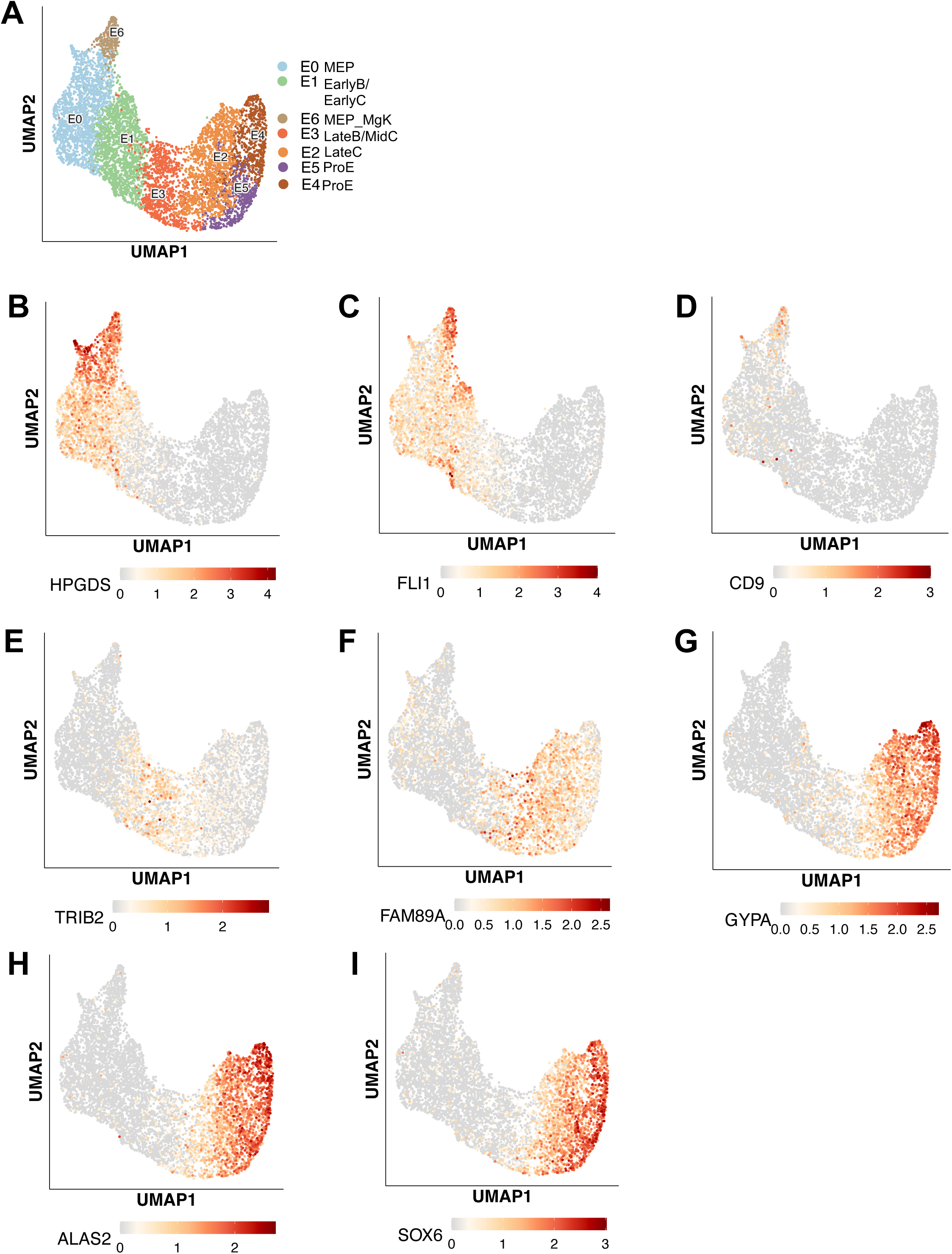
Figure S4.

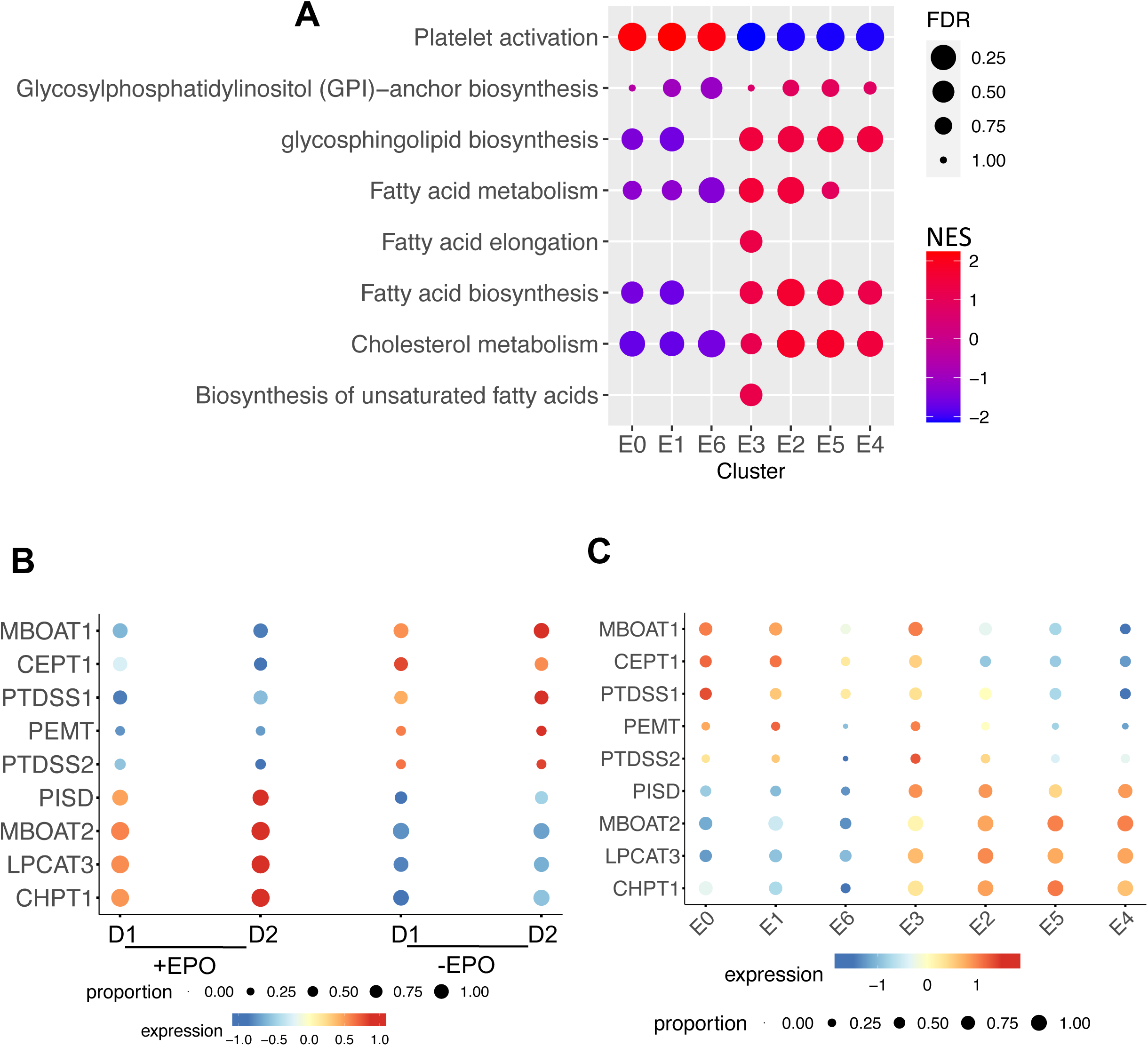
Figure S5.

## References

1 Corrons JLV, Casafont LB, Frasnedo EF. Concise review: how do red blood cells born, live, and die? Ann Hematol 2021;100(10):2425–2433.

2 Cosgrove J, Hustin LSP, de Boer RJ et al. Hematopoiesis in numbers. Trends Immunol 2021;42(12):1100–1112.

3 Lodish H, Flygare J, Chou S. From stem cell to erythroblast: regulation of red cell production at multiple levels by multiple hormones. IUBMB Life 2010;62(7):492–496.

4 Red Cell Membrane Lipids and Aging. Berlin, Heidelberg 1988. Springer Berlin Heidelberg.

5 Bernhardt I, Kaestner L. Historical View and Some Unsolved Problems in Red Blood Cell Membrane Research. Front Biosci (Landmark Ed) 2025;30(3):25331.

6 Mohandas N, Gallagher PG. Red cell membrane: past, present, and future. Blood 2008;112(10):3939–3948.

7 Daleke DL. Regulation of phospholipid asymmetry in the erythrocyte membrane. Curr Opin Hematol 2008;15(3):191–195.

8 Meurs I, Hoekstra M, van Wanrooij EJ et al. HDL cholesterol levels are an important factor for determining the lifespan of erythrocytes. Exp Hematol 2005;33(11):1309–1319.

9 Bernecker C, Kofeler H, Pabst G et al. Cholesterol Deficiency Causes Impaired Osmotic Stability of Cultured Red Blood Cells. Front Physiol 2019;10:1529.

10 Nemkov T, Kingsley PD, Dzieciatkowska M et al. Circulating primitive murine erythroblasts undergo complex proteomic and metabolomic changes during terminal maturation. Blood Adv 2022;6(10):3072–3089.

11 Yamaguchi T, Ishimatu T. Effects of Cholesterol on Membrane Stability of Human Erythrocytes. Biol Pharm Bull 2020;43(10):1604–1608.

12 Lu Z, Huang L, Li Y et al. Fine-Tuning of Cholesterol Homeostasis Controls Erythroid Differentiation. Adv Sci (Weinh) 2022;9(2):e2102669.

13 Mohandas N, Evans E. Mechanical properties of the red cell membrane in relation to molecular structure and genetic defects. Annu Rev Biophys Biomol Struct 1994;23:787–818.

14 Gallagher PG. Chapter 45 – Red Blood Cell Membrane Disorders. In: Hoffman R, Benz EJ, Silberstein LE et al., eds. Hematology (Seventh Edition). Elsevier, 2018:626–647.

15 Kuypers FA. Red cell membrane lipids in hemoglobinopathies. Curr Mol Med 2008;8(7):633–638.

16 Shohet SB, Nathan DG, Livermore BM et al. Hereditary hemolytic anemia associated with abnormal membrane lipid. II. Ion permeability and transport abnormalities. Blood 1973;42(1):1–8.

17 Yawata Y, Kanzaki A, Inoue T et al. Red cell membrane disorders in the Japanese population: clinical, biochemical, electron microscopic, and genetic studies. Int J Hematol 1994;60(1):23–38.

18 Clark MR, Shohet SB, Gottfried EL. Hereditary hemolytic disease with increased red blood cell phosphatidylcholine and dehydration: one, two, or many disorders? Am J Hematol 1993;42(1):25–30.

19 Yawata Y, Kanzaki A, Yawata A et al. Hereditary Red Cell Membrane Disorders in Japan: Their Genotypic and Phenotypic Features in 1014 Cases Studied. Hematology 2001;6(6):399–422.

20 Shohet SB, Livermore BM, Nathan DG et al. Hereditary hemolytic anemia associated with abnormal membrane lipids: mechanism of accumulation of phosphatidyl choline. Blood 1971;38(4):445–456.

21 Yawata Y, Sugihara T, Mori M et al. Lipid analyses and fluidity studies by electron spin resonance of red cell membranes in hereditary high red cell membrane phosphatidylcholine hemolytic anemia. Blood 1984;64(5):1129–1134.

22 Rabelo IB, Chiba AK, Moritz E et al. Metabolomic profile in patients with primary warm autoimmune haemolytic anaemia. Br J Haematol 2023;201(1):140–149.

23 Wiley JS, Ellory JC, Shuman MA et al. Characteristics of the membrane defect in the hereditary stomatocytosis syndrome. Blood 1975;46(3):337–356.

24 Owen JS, Bruckdorfer KR, Day RC et al. Decreased erythrocyte membrane fluidity and altered lipid composition in human liver disease. J Lipid Res 1982;23(1):124–132.

25 Wu H, Bogdanov M, Zhang Y et al. Hypoxia-mediated impaired erythrocyte Lands’ Cycle is pathogenic for sickle cell disease. Sci Rep 2016;6:29637.

26 Peikert K, Spranger A, Miltenberger-Miltenyi G et al. Phosphatidylethanolamines are the Main Lipid Class Altered in Red Blood Cells from Patients with VPS13A Disease/Chorea-Acanthocytosis. Mov Disord 2025;40(3):544–549.

27 Cloos AS, Ghodsi M, Stommen A et al. Red blood cell lipid distribution in the pathophysiology and laboratory evaluation of chorea-acanthocytosis and McLeod syndrome patients. Front Physiol 2025;16:1543812.

28 Allen DW, Manning N. Cholesterol-loading of membranes of normal erythrocytes inhibits phospholipid repair and arachidonoyl-CoA:1-palmitoyl-sn-glycero-3-phosphocholine acyl transferase. A model of spur cell anemia. Blood 1996;87(8):3489–3493.

29 Balaian E, Wobus M, Bornhauser M et al. Myelodysplastic Syndromes and Metabolism. Int J Mol Sci 2021;22(20).

30 Poulaki A, Katsila T, Stergiou IE et al. Bioenergetic Profiling of the Differentiating Human MDS Myeloid Lineage with Low and High Bone Marrow Blast Counts. Cancers (Basel) 2020;12(12).

31 Deng K, Wang Y, Min JC et al. A lipid metabolism defect is an underlying contributor to Diamond Blackfan anemia syndrome. bioRxiv 2025.

32 Mori Y, Chen JY, Pluvinage JV et al. Prospective isolation of human erythroid lineage-committed progenitors. Proc Natl Acad Sci U S A 2015;112(31):9638–9643.

33 Schippel N, Sharma S. Dynamics of human hematopoietic stem and progenitor cell differentiation to the erythroid lineage. Exp Hematol 2023;123:1–17.

34 Doulatov S, Notta F, Laurenti E et al. Hematopoiesis: a human perspective. Cell Stem Cell 2012;10(2):120–136.

35 Constantinescu SN, Ghaffari S, Lodish HF. The Erythropoietin Receptor: Structure, Activation and Intracellular Signal Transduction. Trends Endocrinol Metab 1999;10(1):18–23.

36 Mulcahy L. The erythropoietin receptor. Semin Oncol 2001;28(2 Suppl 8):19–23.

37 Witthuhn BA, Quelle FW, Silvennoinen O et al. JAK2 associates with the erythropoietin receptor and is tyrosine phosphorylated and activated following stimulation with erythropoietin. Cell 1993;74(2):227–236.

38 Bhoopalan SV, Huang LJ, Weiss MJ. Erythropoietin regulation of red blood cell production: from bench to bedside and back. F1000Res 2020;9.

39 Kuhrt D, Wojchowski DM. Emerging EPO and EPO receptor regulators and signal transducers. Blood 2015;125(23):3536–3541.

40 Tothova Z, Semelakova M, Solarova Z et al. The Role of PI3K/AKT and MAPK Signaling Pathways in Erythropoietin Signalization. Int J Mol Sci 2021;22(14).

41 Tothova Z, Tomc J, Debeljak N et al. STAT5 as a Key Protein of Erythropoietin Signalization. Int J Mol Sci 2021;22(13).

42 Tsiftsoglou AS. Erythropoietin (EPO) as a Key Regulator of Erythropoiesis, Bone Remodeling and Endothelial Transdifferentiation of Multipotent Mesenchymal Stem Cells (MSCs): Implications in Regenerative Medicine. Cells 2021;10(8).

43 Li J, Hale J, Bhagia P et al. Isolation and transcriptome analyses of human erythroid progenitors: BFU-E and CFU-E. Blood 2014;124(24):3636–3645.

44 Minetti G, Migliaccio AR, Fibach E. Editorial: Membrane Processes in Erythroid Development and Red Cell Life Time. Front Physiol 2021;12:655117.

45 Minetti G, Achilli C, Perotti C et al. Continuous Change in Membrane and Membrane-Skeleton Organization During Development From Proerythroblast to Senescent Red Blood Cell. Front Physiol 2018;9:286.

46 Minetti G, Dorn I, Kofeler H et al. Insights from lipidomics into the terminal maturation of circulating human reticulocytes. Cell Death Discov 2025;11(1):79.

47 Liao R, Babatunde A, Qiu S et al. A transcriptional network governing ceramide homeostasis establishes a cytokine-dependent developmental process. Nat Commun 2023;14(1):7262.

48 Wu H, Liu X, Jaenisch R et al. Generation of committed erythroid BFU-E and CFU-E progenitors does not require erythropoietin or the erythropoietin receptor. Cell 1995;83(1):59–67.

49 Lin CS, Lim SK, D’Agati V et al. Differential effects of an erythropoietin receptor gene disruption on primitive and definitive erythropoiesis. Genes Dev 1996;10(2):154–164.

50 Kieran MW, Perkins AC, Orkin SH et al. Thrombopoietin rescues in vitro erythroid colony formation from mouse embryos lacking the erythropoietin receptor. Proc Natl Acad Sci U S A 1996;93(17):9126–9131.

51 Malik J, Kim AR, Tyre KA et al. Erythropoietin critically regulates the terminal maturation of murine and human primitive erythroblasts. Haematologica 2013;98(11):1778–1787.

52 Bapat A, Schippel N, Shi X et al. Hypoxia promotes erythroid differentiation through the development of progenitors and proerythroblasts. Exp Hematol 2021;97:32–46 e35.

53 Bapat A, Keita N, Sharma S. Pan-myeloid Differentiation of Human Cord Blood Derived CD34+ Hematopoietic Stem and Progenitor Cells. J Vis Exp 2019(150).

54 Bapat A, Keita N, Martelly W et al. Myeloid Disease Mutations of Splicing Factor SRSF2 Cause G2-M Arrest and Skewed Differentiation of Human Hematopoietic Stem and Progenitor Cells. Stem Cells 2018;36(11):1663–1675.

55 Schippel N, Kala M, Sharma S. Erythropoietin-dependent Acquisition of CD71hiCD105hi Phenotype within CD235a-Early Erythroid Progenitors. Stem Cells 2025.

56 Subramanian A, Tamayo P, Mootha VK et al. Gene set enrichment analysis: a knowledge-based approach for interpreting genome-wide expression profiles. Proc Natl Acad Sci U S A 2005;102(43):15545–15550.

57 Gaud C, B CS, Nguyen A et al. BioPAN: a web-based tool to explore mammalian lipidome metabolic pathways on LIPID MAPS. F1000Res 2021;10:4.

58 Butler A, Hoffman P, Smibert P et al. Integrating single-cell transcriptomic data across different conditions, technologies, and species. Nat Biotechnol 2018;36(5):411–420.

59 Eisele AS, Cosgrove J, Magniez A et al. Erythropoietin directly remodels the clonal composition of murine hematopoietic multipotent progenitor cells. Elife 2022;11.

60 Tusi BK, Wolock SL, Weinreb C et al. Population snapshots predict early haematopoietic and erythroid hierarchies. Nature 2018;555(7694):54–60.

61 Huang P, Zhao Y, Zhong J et al. Putative regulators for the continuum of erythroid differentiation revealed by single-cell transcriptome of human BM and UCB cells. Proc Natl Acad Sci U S A 2020;117(23):12868–12876.

62 Doty RT, Lausted CG, Munday AD et al. The transcriptomic landscape of normal and ineffective erythropoiesis at single-cell resolution. Blood Adv 2023;7(17):4848–4868.

63 An X, Schulz VP, Li J et al. Global transcriptome analyses of human and murine terminal erythroid differentiation. Blood 2014;123(22):3466–3477.

64 Lyu J, Ni M, Weiss MJ et al. Metabolic regulation of erythrocyte development and disorders. Exp Hematol 2024;131:104153.

65 Morita SY, Ikeda Y. Regulation of membrane phospholipid biosynthesis in mammalian cells. Biochem Pharmacol 2022;206:115296.

66 Dorighello G, McPhee M, Halliday K et al. Differential contributions of phosphotransferases CEPT1 and CHPT1 to phosphatidylcholine homeostasis and lipid droplet biogenesis. J Biol Chem 2023;299(4):104578.

67 Hess SY, Owais A, Jefferds MED et al. Accelerating action to reduce anemia: Review of causes and risk factors and related data needs. Ann N Y Acad Sci 2023;1523(1):11–23.

68 Collaborators GBDA. Prevalence, years lived with disability, and trends in anaemia burden by severity and cause, 1990-2021: findings from the Global Burden of Disease Study 2021. Lancet Haematol 2023;10(9):e713–e734.

69 Triana S, Vonficht D, Jopp-Saile L et al. Single-cell proteo-genomic reference maps of the hematopoietic system enable the purification and massive profiling of precisely defined cell states. Nat Immunol 2021;22(12):1577–1589.

70 Zeng AGX, Iacobucci I, Shah S et al. Single-cell Transcriptional Atlas of Human Hematopoiesis Reveals Genetic and Hierarchy-Based Determinants of Aberrant AML Differentiation. Blood Cancer Discov 2025:OF1–OF18.

71 Komic H, Schmachtel T, Simoes C et al. Continuous map of early hematopoietic stem cell differentiation across human lifetime. Nat Commun 2025;16(1):2287.

72 Zhang X, Song B, Carlino MJ et al. An immunophenotype-coupled transcriptomic atlas of human hematopoietic progenitors. Nat Immunol 2024;25(4):703–715.

73 Notta F, Zandi S, Takayama N et al. Distinct routes of lineage development reshape the human blood hierarchy across ontogeny. Science 2016;351(6269):aab2116.

74 Belluschi S, Calderbank EF, Ciaurro V et al. Myelo-lymphoid lineage restriction occurs in the human haematopoietic stem cell compartment before lymphoid-primed multipotent progenitors. Nat Commun 2018;9(1):4100.

75 Pellin D, Loperfido M, Baricordi C et al. A comprehensive single cell transcriptional landscape of human hematopoietic progenitors. Nat Commun 2019;10(1):2395.

76 Gao S, Wu Z, Kannan J et al. Comparative Transcriptomic Analysis of the Hematopoietic System between Human and Mouse by Single Cell RNA Sequencing. Cells 2021;10(5).

77 Giladi A, Paul F, Herzog Y et al. Single-cell characterization of haematopoietic progenitors and their trajectories in homeostasis and perturbed haematopoiesis. Nat Cell Biol 2018;20(7):836–846.

78 Laurenti E, Gottgens B. From haematopoietic stem cells to complex differentiation landscapes. Nature 2018;553(7689):418–426.

79 Zheng S, Papalexi E, Butler A et al. Molecular transitions in early progenitors during human cord blood hematopoiesis. Mol Syst Biol 2018;14(3):e8041.

80 Drissen R, Thongjuea S, Theilgaard-Monch K et al. Identification of two distinct pathways of human myelopoiesis. Sci Immunol 2019;4(35).

81 Doulatov S, Notta F, Eppert K et al. Revised map of the human progenitor hierarchy shows the origin of macrophages and dendritic cells in early lymphoid development. Nat Immunol 2010;11(7):585–593.

82 Jelkmann W. Erythropoietin after a century of research: younger than ever. Eur J Haematol 2007;78(3):183–205.

83 Gao X, Lee HY, da Rocha EL et al. TGF-beta inhibitors stimulate red blood cell production by enhancing self-renewal of BFU-E erythroid progenitors. Blood 2016;128(23):2637–2641.

84 Xie X, Liu M, Zhang Y et al. Single-cell transcriptomic landscape of human blood cells. Natl Sci Rev 2021;8(3):nwaa180.

85 Xu C, He J, Wang H et al. Single-cell transcriptomic analysis identifies an immune-prone population in erythroid precursors during human ontogenesis. Nat Immunol 2022;23(7):1109–1120.

86 Perik-Zavodskii R, Perik-Zavodskaia O, Alrhmoun S et al. Single-cell multi-omics reveal stage of differentiation and trajectory-dependent immunity-related gene expression patterns in human erythroid cells. Front Immunol 2024;15:1431303.

87 Wang B, Tontonoz P. Phospholipid Remodeling in Physiology and Disease. Annu Rev Physiol 2019;81:165–188.

88 Huang NJ, Lin YC, Lin CY et al. Enhanced phosphocholine metabolism is essential for terminal erythropoiesis. Blood 2018;131(26):2955–2966.

89 Oburoglu L, Romano M, Taylor N et al. Metabolic regulation of hematopoietic stem cell commitment and erythroid differentiation. Curr Opin Hematol 2016;23(3):198–205.

90 Ni M, Scaramellini N, Motta I et al. Understanding and targeting erythroid cell metabolism. Trends Cell Biol 2025.

91 D’Alessandro A, Anastasiadi AT, Tzounakas VL et al. Red Blood Cell Metabolism In Vivo and In Vitro. Metabolites 2023;13(7).

92 Orsini M, Chateauvieux S, Rhim J et al. Sphingolipid-mediated inflammatory signaling leading to autophagy inhibition converts erythropoiesis to myelopoiesis in human hematopoietic stem/progenitor cells. Cell Death Differ 2019;26(9):1796–1812.

93 Bhatia H, Hallock JL, Dutta A et al. Short-chain fatty acid-mediated effects on erythropoiesis in primary definitive erythroid cells. Blood 2009;113(25):6440–6448.

94 Zingariello M, Bardelli C, Sancillo L et al. Dexamethasone Predisposes Human Erythroblasts Toward Impaired Lipid Metabolism and Renders Their ex vivo Expansion Highly Dependent on Plasma Lipoproteins. Front Physiol 2019;10:281.

95 Nemkov T, Skinner SC, Nader E et al. Acute Cycling Exercise Induces Changes in Red Blood Cell Deformability and Membrane Lipid Remodeling. Int J Mol Sci 2021;22(2).

