## Supplemental Information for "Erythropoietin Mediates Glycerophospholipid Remodeling During Human Early Erythropoiesis"

**Single Cell Transcriptomic Analysis Reveals Erythropoietin Mediated Remodeling of the Erythroid Lipidome**

### Supplemental Materials and Methods

#### Culture and Differentiation of $CD34^+$ HSPCs

Human bone marrow (BM)-derived $CD34^+$ hematopoietic stem and progenitor cells (HSPCs), all from female donors between the ages 13 to 34 years, were purchased from Ossium Health (San Francisco, CA, USA). For all donor samples, quality control by flow cytometry indicating >90% viability, via staining with 7AAD, and >90% CD34 positivity, using CD45 and CD34 antibody staining, was provided by the supplier and confirmed by in-house immunophenotyping analysis. Furthermore, Ossium provided negative results for HIV-1/2 antibody, HBV surface antigen, HBV core antibody, HCV antibody, HIV/HBV/HCV nucleic acid amplification, cytomegalovirus, and syphilis tests for all donors. MS-5 cells (used as feeder cells for co-culture with $CD34^+$ HSPCs) were maintained in Dulbecco's modified Eagle's medium (DMEM) with high glucose and L-glutamine (Thermo Fisher Scientific, Waltham, MA), 10% fetal bovine serum (FBS; negative for mycoplasma purchased from Omega Scientific, Tarzana, CA), and 1% penicillin-streptomycin (Pen-Strep, Sigma-Aldrich, St. Louis, MO). MS-5 cells were checked annually for mycoplasma contamination and always yielded negative results.

Culture of BM $CD34^+$ HSPCs was performed as previously described [1-4]. Briefly, on day 0, HSPCs were plated at a density of $1 \times 10^5$ cells/well of a 48-well plate in 200 $\mu$L of stimulation medium, containing serum-free X-VIVO 15 (Lonza, Basel, Switzerland), 2 mM L-glutamine, 50 ng/mL stem cell factor (SCF), 50 ng/mL thrombopoietin (TPO), 20 ng/mL interleukin-3 (IL-3), and 50 ng/mL Flt3-Ligand (Flt-3). On day 1, the stimulated $CD34^+$ cells were collected and plated onto MS-5 cells at a density of $10^4$ cells/well of a 48-well plate in 200 $\mu$L of complete medium (DMEM with high glucose and L-glutamine, 10% FBS, and 1% Pen-Strep, supplemented with 50 ng/mL thrombopoietin (TPO) (Miltenyi Biotec, Auburn, CA), 5 ng/mL stem cell factor (SCF), 5 ng/mL interleukin-3 (IL-3), and 5 ng/mL Flt-3 ligand) with or without 4 U/mL erythropoietin (Epo). All cytokines were purchased from BioBasic, Amherst, NY, or StemCell Technologies, Vancouver, BC, Canada. A half-medium change was performed on day 4.

#### Single Cell RNA Sequencing

Cells were collected from human BM $CD34^+$ HSPC cultures grown in complete medium with and without Epo for seven days, washed, and resuspended at a density of $1 \times 10^3$ cells/$\mu$L in ice-cold 1x PBS. Libraries were prepared using the Chromium Next GEM Single Cell 3' Reagent Kit v3.1 (Dual Index; 10x Genomics, Pleasanton, CA) according to the manufacturer's instructions. Briefly, cells were partitioned into gel beads by running them through the Chromium iX controller (10x Genomics, Pleasanton, CA). Full-length cDNA was generated using the 10x Genomics platform in a Bio-Rad C1000 thermal cycler. Gel beads-in-emulsion (GEMs) were cleaned with Dynabeads cleanup mix (see manufacturer's protocol), and cDNA was amplified by PCR and cleaned with solid-phase reversible immobilization (SPRI) beads and then quantified using a Qubit® 2.0 fluorometer (ThermoFisher; Waltham, MA). For the preparation of 3´ gene expression dual index libraries, cDNA fragmentation, end repair, and A-tailing were performed, and then size selection was done using SPRI beads (10x Genomics, Pleasanton, CA). Oligonucleotide adaptors were ligated, dual index TT (set A) primers were incorporated using PCR, and DNA was again quantified on a Qubit® 2.0 fluorometer. Finally, the libraries were sequenced using the Novogene platform (paired-end sequencing, 150 bp reads; PE150).

#### Single Cell Transcriptomic Data Analysis

After sequencing, CellRanger v7.0.1 (10x Genomics) was used for the processing of raw sequencing data (FASTQ files), mapping reads to the human genome version GRCh38-2020-A, and generation of unique molecular identifier (UMI) counts for gene expression via gene-barcode matrix [5]. The software package Seurat version 4.4.0 was used to import data for all four samples, only considering features observed in more than three cells [6]. Additional quality control was performed, including filtering out cells with less than 500 features (low quality or cell fragments), greater than 7500 features, or >10% of reads mapping to the mitochondrial genome. Expression levels were then log normalized to a factor of 10,000. The top 2000 variable features were identified using the *FindVariableFeatures* function. The data was then scaled (*ScaleData* function), and analyzed by principal component analysis (PCA) using the *runPCA* function. Unsupervised clustering was performed by applying the *FindNeighbors* and *FindClusters* functions. UMAPs were then generated using *runUMAP*. Finally, doublets were identified using the DoubletFinder package in R and subsequently filtered out in Seurat [7]. This process was performed for all four samples (Donor 1 and Donor 2 in +/−Epo conditions; resolution 0.5), and then the samples were merged using the *Merge* function in Seurat. The full list of software and packages used, along with their versions, is provided in Supplemental File 1.

**All-Cell UMAP**

Unsupervised clustering was performed on the merged object (resolution 0.4), and ShinyCell and Seurat were used to visualize UMAPs, boxplots, graphs, and heatmaps [8]. Hematopoietic marker genes were then applied to identify clusters specific to HSPCs, granulocyte-monocyte progenitors (GMPs), megakaryocytes, monocytes, granulocytes, erythrocytes, and early lymphoid progenitors, but not mature T and B cells (Figure S1 and Supplemental File 3) [9-11]. The pattern of canonical human HPSC markers *CD34*, *CD38*, and *PTPRC* (CD45RA) indicated that HSPCs were localized to clusters 0, 9, and 10 (Figure S1B). Expression of lineage markers *IL5RA* (CD125) and *CD14* mapped granulocytes and monocytes to clusters 4 and 7, respectively, and *GP1BA* (CD42) mapped megakaryocytes to clusters 11 and 12 (Figure S1C-E) [12-14]. Furthermore, expression of the early lymphoid genes, *HOPX* and *SPINK2*, mapped to clusters 0 and 9, and the GMP-specific genes, *PRTN3* and *MPO*, mapped to cluster 10 (Figures S1F-G) [10,11]. Expression of mature lymphoid markers *CD19* and *CD3D* was observed in a few cells but was not cluster specific and appeared stochastic (Figure S1H). Finally, expression of CD235a identified clusters 6, 3, 1, and 8 as erythroblast-specific (Figure S1I). CD235a$^{+}$ cells represented a large fraction (~34%) and were aligned with cells derived from cultures containing Epo but absent from cultures lacking Epo (Figure S1J).

**Erythroid UMAP**

For development of the erythroid UMAP, clusters 4, 7, 10, 11, and 12 within the all-cell UMAP containing cell types that were not HSPCs and not involved in erythropoiesis, as well as island clusters containing less than 500 cells (13 and 14), were filtered out using the *Subset* function. Although clusters 0 and 9 expressed early lymphoid markers, they also expressed general HSPC markers and so were not filtered out (see above). The remaining cells were re-clustered as described above (resolution 0.5). Pseudotime analysis was performed using Monocle3 [15,16], and DEG analysis was performed using *FindAllMarkers* (Seurat). Bubble plots depicting relative changes in gene expression between samples and clusters were generated in ShinyCell. Gene set enrichment analysis (GSEA) was performed using the WEB-based GEne SeT AnaLysis Toolkit (WebGestalt) from the list of DEGs across each cluster [17].

Trajectory analyses and expression of previously established marker genes were used to manually annotate the HSPC cluster (Figure 1B-E and Supplemental File 3). HSPC surface markers that are commonly employed for immunophenotyping, including *CD34*, *THY1* (CD90), *Flt3* (CD135), *CD38, IL3RA* (CD123), *PTPRC* (CD45RA), and *KIT* (CD117), were all expressed in C0, C1, and C9 (Figure S2). To annotate these clusters more specifically, we evaluated expression of HSC and MPP markers *CD34*, *HLF*, *THY1*, and *AVP*; CMP and GMP markers *CEBPA* and *CEBPE*; and LMPP markers *AFF3* and *SPINK2*, which mapped HSCs and MPPs to cluster C0, CMPs and GMPs to cluster C1, and LMPPs to cluster C9 (Figure 1B, 1E, and S2)

[11,18-21]. Cluster C2, on the other hand, expresses several MEP genes, including *HPGDS*, *FCER1A*, and *GATA2*, but does not express *FLT3* (CD135).

Similarly, BFU-Es, CFU-Es, and erythroblast clusters were annotated using pseudotime ordering and expression of representative marker genes that have been reported in these populations or described as early erythroid genes in previous RNA-seq analyses [10,11,22-26]. BFU-E genes *CSF2RB*, *CASP3*, and *TRIB2* are highly expressed in C3, whereas CFU-E-specific *AHSP* is expressed in C4. Transcripts for three other CFU-E genes, *SLC16A9*, *FAM89A*, and *CNRIP1*, are also high in C4. These patterns suggest that C3 and C4 contain BFU-Es and CFU-Es, respectively. Significant upregulation of *GYPA* (CD235a) suggests the presence of erythroblasts in C4, C10, C6, C8, C5, and C7 (Figure 2C). ProE-specific genes *SREBF1* and *SYNGR1* are high in both C4 and C10, while other ProE genes, ALAS2 and SOX6, arise in C4 but increase in C10, thereby suggesting that ProEs may be distributed between these two clusters. Similarly, examination of BasoE genes, *E2F8*, *E2F7*, *HIST1H1D*, and *HIST1H1E* suggested their distribution between C6 and C8. Genes for PolyE (*E2F2*, *CA2*, *GYPA*, and *NUSAP1*) and OrthoE (*FOXO3*, *TMCC2*, *DCAF12*, and *MKRN1*) were highly expressed in C5 and C7, respectively. The full list of genes used for manual annotation of clusters is provided in Supplemental File 3.

**Early Erythroid UMAP**

For the early erythropoiesis UMAP, a pipeline similar to the one described above was applied. First, cells from C2, C3, C4, and C10 (of the erythroid UMAP) were isolated using the *Subset* function, and the remaining cells were re-clustered again (resolution 0.4). Downstream analyses, including pseudotime ordering and GSEA, were performed as described above. MEPs mapped to clusters E0 and E6 (previously comprising C2) based on expression of *HPGDS*; however, E6 also expressed megakaryocyte markers *FL1* and *CD9*, suggesting E6 may represent a megakaryocyte-primed MEP (MEP_MgK) (Figure 3 and S4A-D) [11,27]. Expression of TRIB2 and FAM89A mapped BFU-Es and CFU-Es to clusters E1 and E3 and E2 and E5, respectively (Figure S4E-F); however, a sharp increase in the expression of *TFRC* (CD71) between E1 and E3 suggested early BFU-E (early B) and early CFU-E (earlyC) mapped to cluster E1 and late BFU-E (lateB) and mid CFU-E (midC) to cluster E3 (Figure 3E) [4]. Furthermore, the highest expression of *ENG* (CD105) in cluster E2 mapped late CFU-Es (lateCs) there (Figure 3F). Erythroblast marker GYPA (CD235a) and ProE markers ALAS2 and SOX6 mapped ProEs to clusters E5 and E4 (Figure S4G-I) [10,25]. For additional details see main manuscript and Supplemental File 3.

**Visualization of Previously Established Genes Involved in Erythropoiesis**

Expression of genes involved in erythropoiesis was assessed in the erythroid UMAP, both by sample (Donor 1 +Epo, Donor 2 +Epo, Donor 1 -Epo, Donor 2 -Epo) and within each of the eleven clusters, ordered in pseudotime (Figure S3). These included genes previously shown to be Epo-dependent, including master erythroid regulators GATA1 [28] and TAL1 [29], as well as ZFP36L2 [30], GAS5 [31], ERH [32], PIM3 [31], CXXC5 [33], TMEM131 [34], LYL1 [31], HES6 [32], EPOR [35], ERFE [36], PIM1 [31,37], XPO7 [34], JAK2 [38], LXN [32], BCL2L12 [37], and APOL1 [34] (Figure S3A-B). We also visualized expression of other genes that were reported to be involved in erythropoiesis but have not been shown to be directly Epo-dependent: TMCC2 [39], KLF1 [40], PHOSPHO1 [41], ZBTB7A [42], LDB1 [43,44], CTCF [45], BCL11A [46,47], RUNX1 [48], GATA2 [49,50], YY1 [51], CD99 [52], and MYC [53].

**Lipid Extraction and Mass Spectrometry Based Analysis**

Cells from BM $CD34^+$ HSPC cultures (four different donors), grown with and without Epo for 7 days, were harvested, counted, and washed with 1x PBS. Washed cells ($1 \times 10^6$ or more per sample) were transferred to a microcentrifuge tube that was pre-washed with 95% n-

Hexane (UN1208 Hexane, H306-1 Thermo Fisher Scientific, Waltham, MA) and resuspended in 200 μL 0.1x PBS (diluted with liquid chromatography-mass spectrometry (LC-MS) grade water (#W64; Thermo Fisher Scientific, Waltham, MA)), 80 μL methanol (MeOH; LC/MS Grade, #A456-4, Thermo Fisher Scientific, Waltham, MA), and 400 μL tert-butyl methyl ether (MTBE; #650560, Sigma Aldrich, St. Louis, MO) and placed on ice. Samples were mixed by vortexing for 10 seconds, incubated for 30 minutes at -20°C, and then sonicated in an ice/water bath for 15 minutes. After sonication, samples were centrifuged at 14,000 rpm for 10 minutes, resulting in the formation of two layers. From the top lipid metabolites containing MTBE layer, a 300 μL aliquot was transferred to a clean hexane-washed microcentrifuge tube and dried using a vacuum centrifuge.

Next, LC-MS/MS analysis was performed on a Vanquish UPLC-Exploris 240 Orbitrap MS system (Thermo Fisher Scientific, Waltham, MA) as previously described [54-56]. Briefly, each sample was run twice—once for positive and once for negative ion modes—using reverse phase chromatography (Waters XSelect HSS T3 column (150 × 2.1 mm, 2.5 μm particle size; Waters Corporation, Milford, MA); Solvent A: 10 mM ammonium acetate in 60% $H_2O$/40% acetonitrile (ACN), Solvent B: 10 mM ammonium acetate in 90% IPA/10% ACN). Solvent B was used for elution, starting at 50% for 3 minutes, increasing to 100% over the span of 12 minutes, maintaining at 100% for 10 minutes, then decreasing back to 50%. Untargeted data from the 100-2000 m/z range was collected using a mass spectrometer equipped with an electrospray ionization (ESI) source. Chemical standards (~800 lipids; Fisher Scientific (Pittsburg, PA), Sigma-Aldrich (Saint Louis, MO), and Avanti Polar Lipids (Alabaster, AL); internal standards: PC (17:0/17:0) and PG (17:0/17:0)) were used to identify peaks from the MS spectra. For MS data extraction, an absolute intensity threshold of 1,000 and a mass accuracy limit of 5 ppm were used. Lipids were then identified and annotated using retention time (RT), exact mass, MS/MS fragmentation pattern, and isotopic pattern (Thermo LipidSearch 4.2 software; peak picking, alignment, and normalization). Additionally, equal volumes from each sample were pooled together and used as quality control (QC) samples. Only signal/peaks with a coefficient of variance (CV) of < 20% between the two QC samples, and those present in > 80% of all samples, were included for analysis (Supplemental File 7 "CV(QC)<20%").

For each sample, the intensities of identified metabolites were divided by the total number of cells, and duplicate metabolites within samples were summed. Statistical analysis of the resulting metabolite abundances after log transformation and normalization was performed in MetaboAnalyst 6.0 [57]. Prediction of lipid reactions and pathways, and the associated genes that may be affected by Epo, was performed using the BioPAN tool within LIPID MAPS [58].

**Immunophenotyping (Intracellular Flow Cytometry)**

For flow cytometry analyses, cells were harvested and stained with fluorescently labeled antibodies as previously described [1-4]. Briefly, cells were washed, counted using the Countess 3-FL Automated Cell Counter (Thermo Fisher Scientific, Waltham, MA), resuspended in 1 ml phosphate-buffered saline (PBS), and incubated with 1 μl of a fixable viability stain (Live-or-Dye™ 405/545 from Biotium) for 30 minutes at room temperature (RT), as per the manufacture's recommendations. All steps from here forward were performed protected from light. Cells were stained with an antibody cocktail including lineage (Lin) markers CD3, CD10, CD11b, CD19, and CD41a, as well as the progenitor marker CD123, all conjugated to PerCP-Cy5.5, CD34-APC-Cy7, CD71-APC, and CD105-BV421 for 20 minutes on ice. Single stained controls were prepared using compensation beads (UltraComp eBeads from Thermo Fisher Scientific, Waltham, MA) as per the manufacturer's recommendations. To proceed with intracellular staining, cells were fixed with 1% paraformaldehyde (PFA) (Sigma Aldrich, St.

Louis, Missouri) for 15 minutes at RT, permeabilized with ice-cold 70% EtOH for 1 hour and 45 minutes on ice, and blocked with blocking buffer (fluorescence-activated cell sorting (FACS) buffer (PBS with 2% FBS and 5 mM EDTA) with 8% goat serum (Thermo Fisher Scientific, Waltham, MA)) for 30 minutes at RT. Finally, cells were stained with unlabeled rabbit-anti-human IgG primary antibodies for an hour at RT, followed by DyLight 488 goat-anti-rabbit secondary antibody for one hour at RT, and resuspended in FACS buffer. All flow cytometry analysis was performed on a Canto II (BD Biosciences, San Jose, CA), and data was analyzed using FlowJo (version 10.9.0; flowjo.com). A complete list of antibody clones with catalog numbers is provided in Supplemental File 2.

**Statistical Analysis**

Four biological replicates were used for untargeted lipidomic analysis, and three or four biological replicates were performed for intracellular flow cytometry validation of GPL enzyme expression. Each biological replicate represented a unique HSPC donor. Graphical representation plots and statistical analysis by student T-test were performed in GraphPad Prism (version 10.0.3; La Jolla, CA). Data is represented as mean ± standard error of the mean (SEM), and p-values $< 0.05$ were considered significant.

### Supplemental Figure Legends

**Figure S1. Single cell transcriptomic analysis of hematopoietic markers. (A)** UMAP visualization of all cells passing quality control from cultures of human BM $CD34^+$ HSPCs analyzed by scRNA-seq. UMAP plots showing expression of markers specific to (**B**) HSPCs, **(C)** granulocytes, **(D)** monocytes, **(E)** megakaryocytes, **(F)** early lymphoid cells, **(G)** GMPs, **(H)** mature lymphoid cells, and **(I)** erythrocytes. Based on these, cluster 0 and 9 were determined to be comprised of HSPCs, cluster 4 of granulocytes, cluster 7 of monocytes, cluster 11 and 12 of megakaryocytes, cluster 10 is of GMPs, clusters 6, 3, 1, and 8 are of erythroid cells, and clusters 13 and 14 as island clusters. Abbreviations: HSPC; hematopoietic stem and progenitor cell, GMP; granulocyte-monocyte progenitor, Epo; erythropoietin.

**Figure S2. Expression patterns of HSPC genes in the erythroid UMAP.** UMAP visualization of cells involved in erythropoiesis colored by **(A)** cluster (for reference; same as in Figure 1B) and **(B)** scaled expression of HSPC and erythroid marker genes. Abbreviations: HSC; hematopoietic stem cell, MPP; multipotent progenitor, CMP; common myeloid progenitor, GMP; granulocyte-monocyte progenitor, LMPP; lympho-myeloid primed progenitor, MEP; megakaryocyte-erythroid progenitor, BFU-E; burst-forming unit erythroid, CFU-E; colony-forming unit erythroid, ProE; proerythroblast, BasoE; basophilic erythroblast, PolyE; polychromatic erythroblast, OrthoE; orthochromatic erythroblast.

**Figure S3. Expression of Epo-associated genes in HSPC and erythroid clusters.** Bubble plots demonstrating gene expression patterns in HSPCs and erythroid cells. **(A)** Genes known to be regulated by Epo are depicted by treatment with (+) and without (-) Epo for each donor. **(B)** Genes known to be regulated by Epo are depicted by cluster in the erythroid UMAP, ordered in pseudotime (Figure 1B). **(C)** Genes previously shown to be involved in erythropoiesis but not directly controlled by Epo are depicted by treatment with (+) and without (-) Epo in each donor. **(D)** Genes previously shown to be involved in erythropoiesis but not directly controlled by Epo are depicted by cluster in the erythroid UMAP, ordered in pseudotime. Abbreviations: Epo; erythropoietin, D1; donor 1, D2; donor 2.

**Figure S4. Annotation of clusters in the early erythroid UMAP.** From the erythroid UMAP, clusters C2 (MEPs), C3 (BFU-E), C4(CFU-E/ProE), and C10 (ProE) were subset and re-clustered, and then manually annotated. **(A)** Early erythroid UMAP showing manual annotation, for reference. UMAPs depicting expression of markers: **(B)** *HPGDS* (MEP marker), **(C)** *FLI1* (early megakaryocyte marker), **(D)** *CD9* (early megakaryocyte marker), **(E)** *TRIB2* (BFU-E marker), **(F)** *FAM89A* (CFU-E marker), **(G)** *GYPA* (erythroblast marker), **(H)** *ALAS2* (ProE marker), and **(I)** *SOX6* (ProE marker). Abbreviations: MEP; megakaryocyte-erythroid progenitor, MgK; megakaryocyte, EarlyB; early BFU-E, LateB; late BFU-E, EarlyC; early CFU-E, MidC, mid CFU-E, LateC; late CFU-E, ProE; proerythroblast.

**Figure S5. Epo-mediated changes in lipid metabolism in early erythropoiesis.** Changes in gene ontology terms and expression of genes involved in glycerophospholipid (GPL) biosynthesis were assessed within the early erythroid UMAP. (**A**) Bubble plot of terms found to be enriched in each cluster by GSEA. The dot color represents the normalized enrichment score (NES), and the size indicates the false discovery rate (FDR), which range from <0.001 to 1. **(B)** Bubble plot depicting relative expression of GPL metabolism genes by donor and treatment, showing Epo-dependence (Figure 6). **(C)** Expression of GPL metabolism genes in clusters of the EE UMAP indicating an Epo-dependent switch. Dot color represents normalized expression, and size indicates the proportion of cells that express the gene within a cluster. Abbreviations: NES; normalized enrichment score, FDR; false discovery rate, Epo; erythropoietin, D1; donor 1,

D2; donor 2, MBOAT(1 or 2); membrane bound glycerophospholipid O-acyltransferase (1 or 2), CEPT1; choline/ethanolamine phosphotransferase 1, PTDSS(1 or 2); phosphatidylserine synthase (1 or 2), PEMT; phosphatidylethanolamine N-methyltransferase, PISD; phosphatidylserine decarboxylase, LPCAT3; lysophosphatidylcholine acyltransferase 3, CHPT1; Choline phosphotransferase 1.

### References


1 Bapat A, Schippel N, Shi X et al. Hypoxia promotes erythroid differentiation through the development of progenitors and proerythroblasts. Exp Hematol 2021;97:32–46 e35.

2 Bapat A, Keita N, Sharma S. Pan-myeloid Differentiation of Human Cord Blood Derived CD34+ Hematopoietic Stem and Progenitor Cells. J Vis Exp 2019(150).

3 Bapat A, Keita N, Martelly W et al. Myeloid Disease Mutations of Splicing Factor SRSF2 Cause G2-M Arrest and Skewed Differentiation of Human Hematopoietic Stem and Progenitor Cells. Stem Cells 2018;36(11):1663–1675.

4 Schippel N, Kala M, Sharma S. Erythropoietin-dependent Acquisition of CD71hiCD105hi Phenotype within CD235a- Early Erythroid Progenitors. Stem Cells 2025.

5 Zheng GX, Terry JM, Belgrader P et al. Massively parallel digital transcriptional profiling of single cells. Nat Commun 2017;8:14049.

6 Hao Y, Hao S, Andersen-Nissen E et al. Integrated analysis of multimodal single-cell data. Cell 2021;184(13):3573–3587 e3529.

7 McGinnis CS, Murrow LM, Gartner ZJ. DoubletFinder: Doublet Detection in Single-Cell RNA Sequencing Data Using Artificial Nearest Neighbors. Cell Syst 2019;8(4):329–337 e324.

8 Ouyang JF, Kamaraj US, Cao EY et al. ShinyCell: simple and sharable visualization of single-cell gene expression data. Bioinformatics 2021;37(19):3374–3376.

9 Notta F, Zandi S, Takayama N et al. Distinct routes of lineage development reshape the human blood hierarchy across ontogeny. Science 2016;351(6269):aab2116.

10 Zeng AGX, Iacobucci I, Shah S et al. Single-cell Transcriptional Atlas of Human Hematopoiesis Reveals Genetic and Hierarchy-Based Determinants of Aberrant AML Differentiation. Blood Cancer Discov 2025:OF1–OF18.

11 Ainciburu M, Ezponda T, Berastegui N et al. Uncovering perturbations in human hematopoiesis associated with healthy aging and myeloid malignancies at single-cell resolution. Elife 2023;12.

12 Friedman AD. Transcriptional regulation of granulocyte and monocyte development. Oncogene 2002;21(21):3377–3390.

13 Papalexi E, Satija R. Single-cell RNA sequencing to explore immune cell heterogeneity. Nat Rev Immunol 2018;18(1):35–45.

14 Zhang W, Yan C, Liu X et al. Global characterization of megakaryocytes in bone marrow, peripheral blood, and cord blood by single-cell RNA sequencing. Cancer Gene Ther 2022;29(11):1636–1647.

15 Trapnell C, Cacchiarelli D, Grimsby J et al. The dynamics and regulators of cell fate decisions are revealed by pseudotemporal ordering of single cells. Nat Biotechnol 2014;32(4):381–386.

16 Cao J, Spielmann M, Qiu X et al. The single-cell transcriptional landscape of mammalian organogenesis. Nature 2019;566(7745):496–502.

17 Elizarraras JM, Liao Y, Shi Z et al. WebGestalt 2024: faster gene set analysis and new support for metabolomics and multi-omics. Nucleic Acids Res 2024;52(W1):W415–W421.

18 Komic H, Schmachtel T, Simoes C et al. Continuous map of early hematopoietic stem cell differentiation across human lifetime. Nat Commun 2025;16(1):2287.

19 Zhang X, Song B, Carlino MJ et al. An immunophenotype-coupled transcriptomic atlas of human hematopoietic progenitors. Nat Immunol 2024;25(4):703–715.

20 Gao S, Wu Z, Kannan J et al. Comparative Transcriptomic Analysis of the Hematopoietic System between Human and Mouse by Single Cell RNA Sequencing. Cells 2021;10(5).

21 Kucinski I, Campos J, Barile M et al. A time- and single-cell-resolved model of murine bone marrow hematopoiesis. Cell Stem Cell 2024;31(2):244–259 e210.

22 Li J, Hale J, Bhagia P et al. Isolation and transcriptome analyses of human erythroid progenitors: BFU-E and CFU-E. Blood 2014;124(24):3636–3645.
23 Huang P, Zhao Y, Zhong J et al. Putative regulators for the continuum of erythroid differentiation revealed by single-cell transcriptome of human BM and UCB cells. Proc Natl Acad Sci U S A 2020;117(23):12868–12876.
24 Pellin D, Loperfido M, Baricordi C et al. A comprehensive single cell transcriptional landscape of human hematopoietic progenitors. Nat Commun 2019;10(1):2395.
25 Dumitriu B, Patrick MR, Petschek JP et al. Sox6 cell-autonomously stimulates erythroid cell survival, proliferation, and terminal maturation and is thereby an important enhancer of definitive erythropoiesis during mouse development. Blood 2006;108(4):1198–1207.
26 An X, Schulz VP, Li J et al. Global transcriptome analyses of human and murine terminal erythroid differentiation. Blood 2014;123(22):3466–3477.
27 Psaila B, Barkas N, Iskander D et al. Single-cell profiling of human megakaryocyte-erythroid progenitors identifies distinct megakaryocyte and erythroid differentiation pathways. Genome Biol 2016;17:83.
28 Ferreira R, Ohneda K, Yamamoto M et al. GATA1 function, a paradigm for transcription factors in hematopoiesis. Mol Cell Biol 2005;25(4):1215–1227.
29 Prasad KS, Jordan JE, Koury MJ et al. Erythropoietin stimulates transcription of the TAL1/SCL gene and phosphorylation of its protein products. J Biol Chem 1995;270(19):11603–11611.
30 Zhang L, Prak L, Rayon-Estrada V et al. ZFP36L2 is required for self-renewal of early burst-forming unit erythroid progenitors. Nature 2013;499(7456):92–96.
31 Singh S, Dev A, Verma R et al. Defining an EPOR- regulated transcriptome for primary progenitors, including Tnfr-sf13c as a novel mediator of EPO- dependent erythroblast formation. PLoS One 2012;7(7):e38530.
32 da Cunha AF, Brugnerotto AF, Duarte AS et al. Global gene expression reveals a set of new genes involved in the modification of cells during erythroid differentiation. Cell Prolif 2010;43(3):297–309.
33 Astori A, Matherat G, Munoz I et al. The epigenetic regulator RINF (CXXC5) maintains SMAD7 expression in human immature erythroid cells and sustains red blood cells expansion. Haematologica 2022;107(1):268–283.
34 Koury S, Yarlagadda S, Moskalik-Liermo K et al. Differential gene expression during terminal erythroid differentiation. Genomics 2007;90(5):574–582.
35 Chiba T, Ikawa Y, Todokoro K. GATA-1 transactivates erythropoietin receptor gene, and erythropoietin receptor-mediated signals enhance GATA-1 gene expression. Nucleic Acids Res 1991;19(14):3843–3848.
36 Kautz L, Jung G, Valore EV et al. Identification of erythroferrone as an erythroid regulator of iron metabolism. Nat Genet 2014;46(7):678–684.
37 Gillinder KR, Tuckey H, Bell CC et al. Direct targets of pSTAT5 signalling in erythropoiesis. PLoS One 2017;12(7):e0180922.
38 Miura O, Miura Y, Nakamura N et al. Induction of tyrosine phosphorylation of Vav and expression of Pim-1 correlates with Jak2-mediated growth signaling from the erythropoietin receptor. Blood 1994;84(12):4135–4141.
39 Ludwig LS, Lareau CA, Bao EL et al. Transcriptional States and Chromatin Accessibility Underlying Human Erythropoiesis. Cell Rep 2019;27(11):3228–3240 e3227.
40 Yien YY, Bieker JJ. EKLF/KLF1, a tissue-restricted integrator of transcriptional control, chromatin remodeling, and lineage determination. Mol Cell Biol 2013;33(1):4–13.
41 Huang NJ, Lin YC, Lin CY et al. Enhanced phosphocholine metabolism is essential for terminal erythropoiesis. Blood 2018;131(26):2955–2966.
42 Norton LJ, Funnell APW, Burdach J et al. KLF1 directly activates expression of the novel fetal globin repressor ZBTB7A/LRF in erythroid cells. Blood Adv 2017;1(11):685–692.

43 Li L, Lee JY, Gross J et al. A requirement for Lim domain binding protein 1 in erythropoiesis. J Exp Med 2010;207(12):2543–2550.
44 Wadman IA, Osada H, Grutz GG et al. The LIM-only protein Lmo2 is a bridging molecule assembling an erythroid, DNA-binding complex which includes the TAL1, E47, GATA-1 and Ldb1/NLI proteins. EMBO J 1997;16(11):3145–3157.
45 Yang X, Cheng L, Xin Y et al. CTCF is selectively required for maintaining chromatin accessibility and gene expression in human erythropoiesis. Genome Biol 2025;26(1):44.
46 Esteghamat F, Gillemans N, Bilic I et al. Erythropoiesis and globin switching in compound Klf1::Bcl11a mutant mice. Blood 2013;121(13):2553–2562.
47 Zheng G, Orkin SH. Transcriptional Repressor BCL11A in Erythroid Cells. Adv Exp Med Biol 2024;1459:199–215.
48 Kuvardina ON, Herglotz J, Kolodziej S et al. RUNX1 represses the erythroid gene expression program during megakaryocytic differentiation. Blood 2015;125(23):3570–3579.
49 Deindl P, Klar M, Drews D et al. Mice over-expressing human erythropoietin indicate that erythropoietin enhances expression of its receptor via up-regulated Gata1 and Tal1. Haematologica 2014;99(10):e205–207.
50 Bresnick EH, Lee HY, Fujiwara T et al. GATA switches as developmental drivers. J Biol Chem 2010;285(41):31087–31093.
51 Perreault AA, Brown JD, Venters BJ. Erythropoietin Regulates Transcription and YY1 Dynamics in a Pre-established Chromatin Architecture. iScience 2020;23(10):101583.
52 Wang X, Zhang W, Zhao S et al. Decoding human in vitro terminal erythropoiesis originating from umbilical cord blood mononuclear cells and pluripotent stem cells. Cell Prolif 2024;57(7):e13614.
53 Jayapal SR, Lee KL, Ji P et al. Down-regulation of Myc is essential for terminal erythroid maturation. J Biol Chem 2010;285(51):40252–40265.
54 Scieszka DP, Garland D, Hunter R et al. Multi-omic assessment shows dysregulation of pulmonary and systemic immunity to e-cigarette exposure. Respir Res 2023;24(1):138.
55 Mi Y, Qi G, Vitali F et al. Loss of fatty acid degradation by astrocytic mitochondria triggers neuroinflammation and neurodegeneration. Nat Metab 2023;5(3):445–465.
56 Eghlimi R, Shi X, Hrovat J et al. Triple Negative Breast Cancer Detection Using LC-MS/MS Lipidomic Profiling. J Proteome Res 2020;19(6):2367–2378.
57 Pang Z, Lu Y, Zhou G et al. MetaboAnalyst 6.0: towards a unified platform for metabolomics data processing, analysis and interpretation. Nucleic Acids Res 2024;52(W1):W398–W406.
58 Gaud C, B CS, Nguyen A et al. BioPAN: a web-based tool to explore mammalian lipidome metabolic pathways on LIPID MAPS. F1000Res 2021;10:4.
